# A long non-coding RNA regulates in vitro and in vivo triazole antifungal susceptibility in *Aspergillus fumigatus*

**DOI:** 10.1101/2025.10.16.682787

**Authors:** Nava Raj Poudyal, Ryan T. Mehlem, Priyanka Doneparthi, Tabitha L. Cady, Garrett Kaufman, Sven D. Willger, Maximiliano Ortiz, Rooksana E. Noorai, Jason E. Stajich, Sourabh Dhingra

**Affiliations:** Department of Biological Sciences, Clemson University, Clemson, SC 29634, USA; Eukaryotic Pathogen Innovation Center, Clemson University, Clemson, SC 29634, USA; Department of Genetics and Biochemistry, Clemson University, Clemson, SC 29634, USA; USC School of Medicine-Greenville, Greenville, SC 29605, USA; Department of Immunology and Infectious Disease, Montana State University, Bozeman, MT 59717, USA; Clemson University Genomics & Bioinformatics Facility, Clemson University, Clemson, SC 29634, USA; Department of Microbiology and Plant Pathology, University of California-Riverside, Riverside, CA 92521, USA

## Abstract

Azole-resistant *Aspergillus* infections are a source of increasing concern with limited alternative therapeutic options. However, as most infections are still caused by azole-susceptible *Aspergillus* strains, there is a need to better understand fungal responses to azole antifungals. To this end, we discover that a long non-coding RNA, *afu-182,* is a major regulator of *cyp51-*independent sub-MIC azole response. We observe that loss of *afu-182* leads to increased surface attached growth and poor treated disease outcomes in a murine model of invasive pulmonary aspergillosis upon azole treatment. In contrast, overexpression of *afu-182* significantly reduces fungal burden in animals treated with the azole drug, posaconazole. Importantly, *afu-182* levels decrease upon azole exposure and in an azole adaptation experiment, continuous exposure to low dose azole led to MIC increase in an *afu-182* dependent manner. Whole transcriptome analyses revealed that azole drug treatment leads to an increase in transcripts of genes encoding 7-transmembrane domain proteins of the RTA1 family, and these proteins are negatively regulated by *afu-182*. Two RTA1 family genes have individual and combined effects and are sufficient to increase fungal susceptibility to azole drugs in the WT strain. Taken together, our data show a role of the long non-coding RNA *afu-182* in regulating *Aspergillus fumigatus* response to azole drugs both *in vitro* and *in vivo*.

**Importance:** Drug resistance in *Aspergillus* is a major challenge that is often associated with the agricultural use of azoles in the environment. How drug resistance arises *in vivo* is still an active area of research. Here, we show that azole exposure results in fungal adaptation by lowering the RNA levels of lncRNA, which upon low dose azole exposure leads to an increase in azole drug MIC.

## Introduction

*Aspergillus fumigatus* causes a spectrum of infections termed as Aspergillosis, mainly in patients with compromised immune systems (1). The patient population most susceptible to infections is those who have acute neutropenia due to underlying conditions or immunotherapies, or patients on long-term steroid use (2, 3). Recent studies have shown that *A. fumigatus* is a significant cause of secondary infections, leading to increased morbidity and mortality in patients with viral infections, including COVID-19 and influenza A virus (4, 5). Importantly, *A. fumigatus* is increasingly a source of infection in tertiary care centers in patients with mild to no immune suppression, highlighting the prevalence of the infection and increase in the patient population that is susceptible to aspergillosis (6). The World Health Organization (WHO) classified drug resistant *A*. *fumigatus* as a critical pathogen in 2023 (7).

Azoles, polyenes, and echinocandins are the current classes of antifungal drugs available for aspergillosis treatment; azoles are the most widely used (8). Clinically, voriconazole is the frontline therapy; however, posaconazole, itraconazole and isavuconazole are also used (8). Concerningly, azole drug resistance is on the rise (9). Azoles inhibit the sterol 14-α-demethylase gene, *cyp51* (*erg11* in yeast), which leads to accumulation of toxic sterols (10). Molecularly, resistance to azoles is primarily attributed to mutational hotspots in the *cyp51* gene sequence that affect drug-enzyme binding (11); with or without the replication of a 34 base pair or 46 base pair tandem repeat (TR) in the promoter region of the *cyp51a* gene (commonly referred to as TR_34_ and TR_46,_ respectively) (12–14). TRs in promoter regions increase the binding site of transcription factor SrbA, which positively regulates *cyp51a* transcription (15). SrbA also regulates ergosterol biosynthesis by transcriptional regulation of *erg3*, *erg25* and *erg24* genes (16, 17). In addition, transcription factor *atrR* cooperates with SrbA for azole resistance (18).

SrbA is a class of ER resident sterol regulator binding element protein (SREBP) that, in association with SREBP cleavage activating protein (Scap) and Insulin induced gene (INSIG), regulates cholesterol levels in mammalian cells (19, 20). In *A*. *fumigatus*, SrbA plays a significant role in the hypoxia response, low iron response, virulence, and azole susceptibility (19); however, a Scap homolog has not been identified in any Eurotiomycete fungus to date, and the Insig homolog *insA* does not regulate SREBPs or subsequent hypoxia response in these fungi (21). In the mammalian system, INSIG senses the accumulation of sterol pathway intermediates and negatively regulates HMG-CoA reductase, thus blocking the rate-limiting step in sterol biosynthesis (22). In *A. fumigatus,* mutations in HMG-CoA reductase (HMG-CoA) have been recently identified in clinical isolates (23–26), and these mutations, especially in the sterol-sensing domain (SSD), are predicted to inhibit sterol intermediate sensing by changing HMG-CoA-Insig stochiometric balance (27). Thus, *insA* provides a feedback mechanism that contributes to azole resistance (27). Interestingly, overexpression (OE) of *insA* reverses the azole resistance in OE HMG-CoA mutants but not in SSD mutants, highlighting the importance of sterol sensing by HMG-CoA (27).

In a laboratory setting, drug resistance is defined as a change in minimum inhibitory concentration (MIC) above the clinical cut-off value measured in standardized MIC testing assays (28). Multiple methods are used to determine the MICs of the antifungal drugs in the laboratory setting, including broth microdilution, disk diffusion assay and E-Test strips (28, 29), and strains are then classified as susceptible, resistant, tolerant, persistent or exhibiting hetero-resistance (30) based on the ability of all cells or a sub-population of cells to grow at or above MIC (30, 31).

Here, we show that *insA* adjacent long non-coding RNA *afu-182* negatively regulates and aids in fungal adaptation to the azole drugs at sub-MIC concentrations. In an animal model of IPA, the Δ*afu-182* strain causes azole recalcitrant infections, whereas OE-182 strains show increased clearance from the lungs when treated with the azole drug, posaconazole. Upon low dose exposure to azoles, Δ*afu-182* strain adapts and has an increase in the MIC for voriconazole, whereas the WT strain does not show a change in MIC.

Collectively, we observed that *afu-182* regulates fungal azole response independent of *cyp51* regulation and plays a significant role in azole treatment outcomes, especially in strains classified as azole-susceptible. Thus, our study has uncovered a novel role of long ncRNA *afu-182* in azole susceptibility both *in vitro* and *in vivo*.

## Results

### *afu-182* is an intergenic long non-coding RNA

*afu-182* was previously characterized and annotated as an anti-sense ncRNA to the gene *insA* (32, 33). To determine if *afu-182* is an anti-sense or intergenic ncRNA, we performed 3’ rapid amplification of cDNA ends (RACE) (34) for *insA* and showed that *insA* transcript ends 191bp after the translational stop codon (Figure 1). A non-species specific computational analysis assigned *afu-182* sequence a Fickett score of 0.40586 with a protein coding probability of 0.016 (35), and is classified as an intergenic long non-coding RNA.

**Figure 1.**
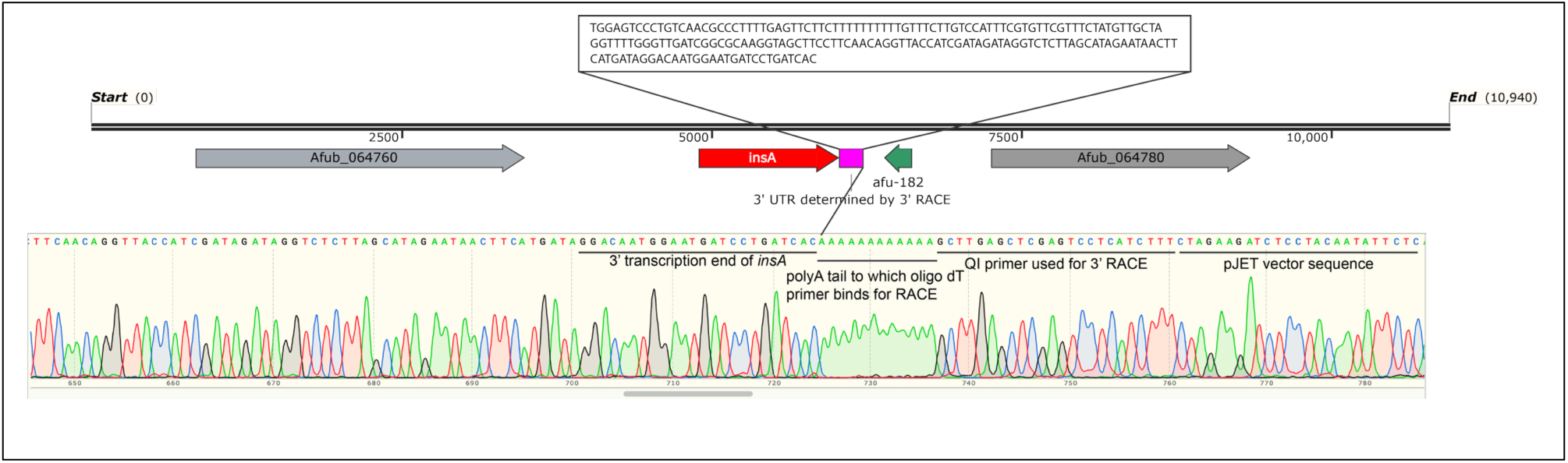
*insA* and *afu-182* are independent transcripts. Schematic showing the genomic location of *afu-182* with respect to *insA* gene. The 3’ end of *insA* was determined via 3’ RACE. The inset sequence shows the sequencing data matching the end of *insA* transcript.

### *afu-182* regulates sub-MIC response to azole class of anti-fungal drugs

*afu-182* neighboring INSIG homolog gene *insA,* was recently shown to play a role in azole drug susceptibility (27) and Supplementary Figure 1A. However, when we complemented *insA* ectopically, we did not see phenotypic reversion (Supplementary Figure 1A, red arrows); even when the *insA* RNA levels were rescued (Supplementary Figure 1B). The Δ*insA* strain does have inherent *afu-182* RNA (Supplementary Figure 1C); however, to determine the cis-acting roles of ncRNA, we also complemented the Δ*insA* strain with DNA fragment containing both *insA* fragment and *afu-182* fragment and saw partial reversal of phenotype (Supplementary Figure 1A, orange arrows); however, no change in voriconazole MIC was observed (Supplementary Figure 1D).

We compared the RNA levels of *afu-182* in Δ*insA* and comp-*insA* strains. Even though, statistically insignificant, we did see 2-fold lower levels of *afu-182* RNA levels in *comp-insA* strain that did not revert the phenotype (Supplementary Figure 1C). Thus, to determine if *afu-182* levels regulate fungal response to sub-MIC azole concentrations, we made a Δ*afu-182* and over-expression 182 (OE-182) strains. We also complemented *afu-182* back at the original locus (Comp-182). We confirmed the levels of *afu-182* in the complement and OE strain and the absence of RNA in the Δ*afu-182* strain (Supplementary Figure 2A). Our qPCR data show the absence of *afu-182* transcript in the Δ*afu-182* strain and WT level of expression in the Comp-182 strain (p=0.5469, One-Way ANOVA, Supplementary Figure 2A). We observed a 4-fold overexpression in the OE-182 strain (p = 0.0025, One-Way ANOVA, Supplementary Figure 2A). To assess the role of *afu-182* in fungal azole response, we serially inoculated the strains (10^5^-10^2^ conidial density) in the absence and presence of 0.25μg/ml voriconazole, itraconazole (0.075μg/ml), posaconazole (0.03μg/ml), and isavuconazole (0.25μg/ml). The *afu-182* null mutant strain showed increased diametric growth against all azoles, confirming that *afu-182* lncRNA regulates sub-MIC fungal azole response to azole class of drugs (Figures 2A – 2E), and importantly this effect was not due to a change in MIC (Supplementary Figure 2B). OE-182 showed decreased diametric fungal growth to azoles, suggesting a causal link between *afu-182 levels* and fungal azole response (Figures 2A-2E).

**Figure 2.**
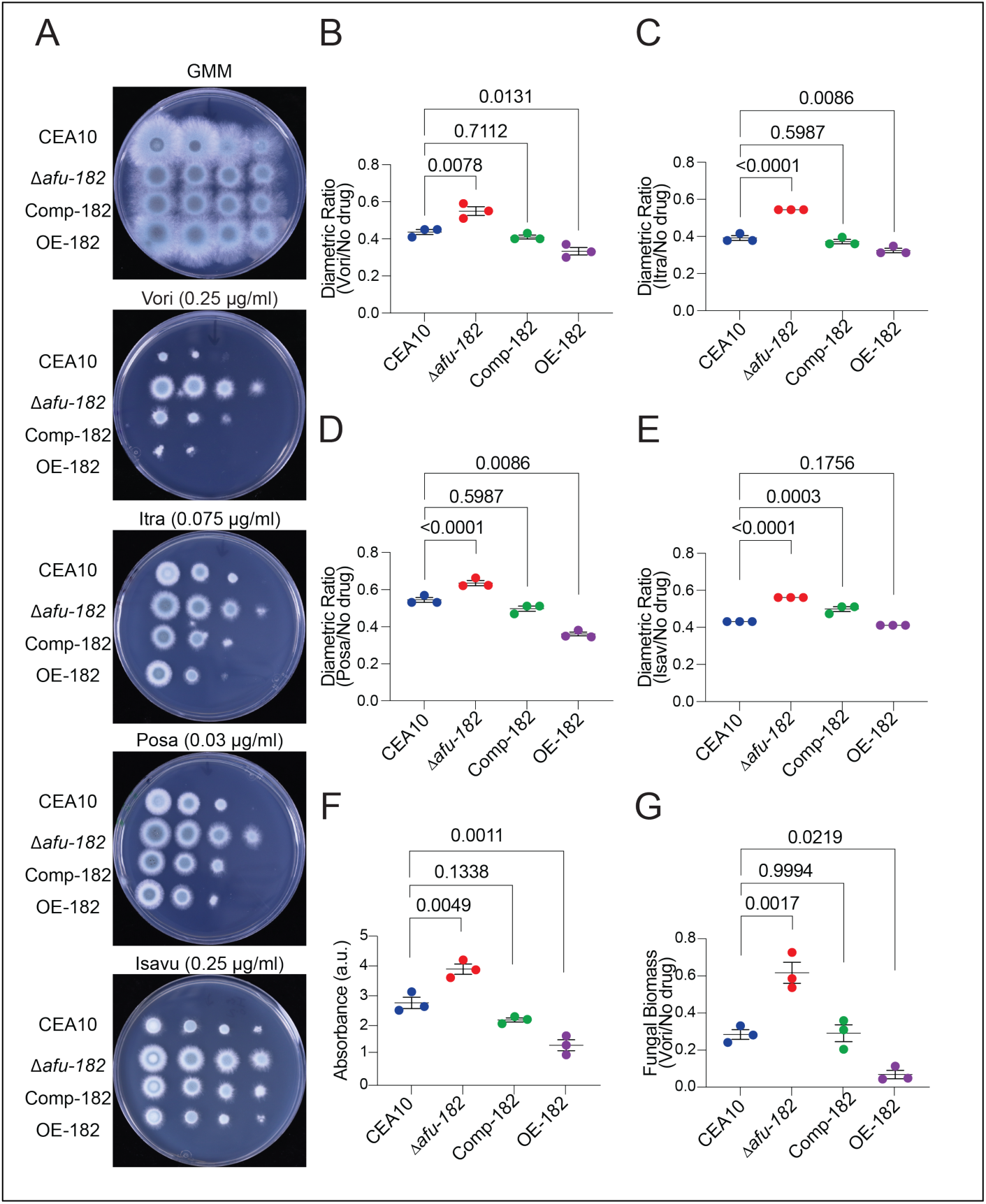
*afu-182* regulates sub-MIC fungal response to azoles. (A) AF strains were serially inoculated (10^5^ – 10^2^) on GMM in the absence and presence of azole drugs as indicated. The Δ*afu-182* strain showed increased fungal growth in the presence of the sub-MIC concentration of voriconazole, whereas the OE-182 strain is more inhibited compared to the WT strain. (B-E) Growth ratio. Colony diameters were measured for colonies as grown in A (for 10^4^ spore density) and represented as diametric growth ratio (Drug/no drug). (F) *afu-182* plays a role in fungal biofilm response to azole drugs. AF strains were allowed to adhere to cell culture plates, and biofilms were treated with voriconazole and stained with crystal violet. The OD_600_ values are represented. Δ*afu-182* formed azole recalcitrant biofilms (p=0.0049, One-Way ANOVA), whereas OE-182 showed a significant decrease in biofilm formation (p=0.0011, One-Way ANOVA) when treated with voriconazole. (G) AF strains were grown in liquid submerged culture with continuous shaking in the absence or presence of 0.2μg/ml voriconazole. After 48 hours, fungal biomass was collected and lyophilized. Dry weight ratio (Drug/no drug) is represented. One-way ANOVA was used to assess the differences in mean; p-values are listed for all samples compared to WT (One-Way ANOVA followed by Tukey post-hoc analysis). a.u. – arbitrary units.

It is possible that *afu-182* inhibits fungal germination and not the azole stress response. Thus, we assessed the role of *afu-182* in fungal azole response post-germination in surface attached cultures. We allowed all strains to germinate and adhere to the abiotic surface (i.e., cell culture plates) for 12 hours. We then washed the fungal biofilms with PBS and grew the biofilms in the presence of voriconazole (0.2μg/ml) for 8 additional hours. The final biofilms were washed, stained with crystal violet and de-stained with ethanol as described previously (36). The binding of crystal violet was measured as an indirect measure of fungal biomass. We observed increased biofilm formation in the Δ*afu-182* upon azole treatment (41% increase compared to WT, p=0.0049, One-Way ANOVA), whereas the OE-182 mutant showed severe and significant impairment in biofilm development after drug treatment (51% decrease compared to WT, p=0.0011, One-Way ANOVA) (Figure 2F), indicating *afu-182* mediated fungal azole response is independent of spore germination. No difference across strains was observed in initial biofilm formation after 12 hours (Supplementary Figure 2C).

Additionally, to understand the role of growth condition-specific role of *afu-182*, we grew AF as liquid submerged cultures in the presence and absence of voriconazole for 48 hours and weighed dry fungal biomass. The Δ*afu-182* strain showed significantly increased biomass (116% increase compared to WT, p=0.0017, One-Way ANOVA), whereas the OE-182 strain showed significant impairment (76% decrease compared to WT, p=0.0219, One-Way ANOVA, Figure 2G). Thus, our data show that *afu-182* influences fungal azole response in both submerged and surface-attached cultures.

### RNA levels of *afu-182* are negatively regulated by azole drugs, and correlate with azole MIC

To understand the role of *afu-182* in fungal azole response, we tested the RNA levels of *afu-182* in the presence and absence of azoles. A significant reduction in levels of *afu-182* was observed in strains grown in the presence of either voriconazole or posaconazole compared to GMM (Figure 3A). Multiple strains of *A. fumigatus,* including A1163, CEA10 and Af293, are used as laboratory WT strains (37). Additionally, multiple reports have shown significant strain virulence heterogeneity, thus making it hard to establish a true WT strain (38); however, a comprehensive study of azole response in ASAF strains is lacking. To further determine the role of *afu-182* in regulating fungal azole response, we quantified the relative abundance (to CEA10) of *afu-182* RNA levels in strain Af293, environmental isolates 47-57, 47-7, 08-19-02-46 (referred as - 46) and 08-19-02-10 (referred as -10). Interestingly, *afu-182* RNA levels differ in AF strains and negatively correlate with azole response i.e. we observed lower MIC for strains trending higher for *afu-182* levels compared to CEA10 (47-7 and 47-57) and increased MIC for strains trending lower for *afu-182* RNA levels compared to CEA10 (Figures 3B and 3C). To confirm if *afu-182* is sufficient to regulate the azole response in multiple isolates, we made a Δ*afu-182* mutant in the Af293.1 background, and the resulting strain showed increased fungal growth at sub-MIC voriconazole concentration of 0.4μg/ml (Figures 3D and 3E). Environmental isolate 47-57 has increased RNA levels of *afu-182* compared to CEA10 and is more susceptible to voriconazole compared to CEA10 (Figure 3C).

**Figure 3.**
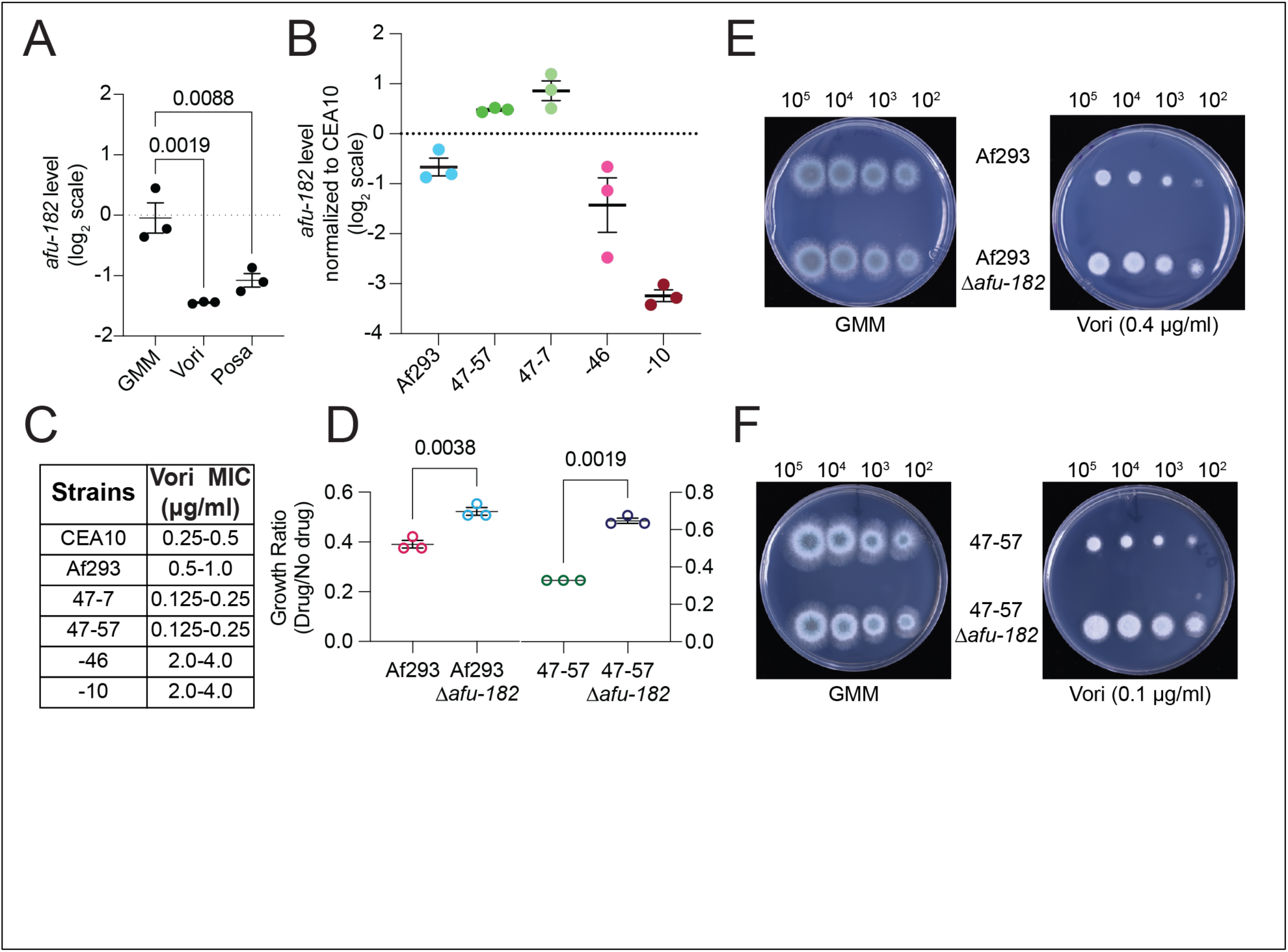
*afu-182* RNA levels are negatively regulated by azoles and correlate with MIC. (A) Quantitative reverse transcriptase PCR was used to determine the *afu-182* RNA levels in the WT strain grown in the absence and presence of azoles. Significant downregulation of *afu-182* was observed in the presence of voriconazole (p=0.0019, One-Way ANOVA) and posaconazole (p=0.0088, One-Way ANOVA). (B) Quantitative reverse transcriptase PCR was used to determine the *afu-182* RNA levels in Af293, 47-57, 47-7, -46 and -10 isolates compared to CEA10. (C) Minimum inhibitory concentration. Voriconazole MIC as measured by broth microdilution method for strains in B. (D) Growth ratio. *afu-182* was deleted from laboratory reference strain Af293.1 and environmental isolate 47-57. WT and corresponding deletion strains were serially inoculated on GMM in the absence and presence of voriconazole. Growth ratio represented as the ratio of colony diameter in the presence of drug over no drug is represented. Significantly more growth was observed in Δ*afu-182* strains for Af293 and 47-57 strains (two-tailed t-test, p =0.0038 and 0.0019, respectively). AF strains were serially inoculated (10^5^ – 10^2^) on GMM in the absence and presence of voriconazole as indicated. The Δ*afu-182* strain showed increased fungal growth in the presence of the sub-MIC concentration of voriconazole.

Deletion of *afu-182* in the 47-57 background also resulted in increased fungal growth of Δ*afu-182* strain compared to the host strain at sub-MIC voriconazole concentration of 0.1μg/ml (Figure 3F). These two isolates with azole sensitivity on either side of CEA10 showed increased fungal growth upon *afu-182* loss, highlighting the role of *afu-182* irrespective of the strain’s background. To further characterize the role of *afu-182* in fungal azole response, we identified the *afu-182* homolog in the related species *A*. *nidulans*. The deletion of *an-182* (Δ*an-182*) in two distinct AN strains (A1145 and GR5) also leads to increased fungal growth (Supplementary Figure 3, orange arrows), indicating roles of lncRNA *afu-*182 homologs in multiple Aspergilli species.

### *afu-182* does not regulate transcription of azole target gene *cyp51* or transcription factors *srbA* and *atrR*

To further test the role *of afu-182* in regulating fugal sub-MIC azole response, we quantified the mRNA levels of azole target genes *cyp51a* and *cyp51b,* both in the presence and absence of azoles and TFs *srbA* and *atrR* in GMM. Importantly, we did not see a change in the transcript levels of *cyp51* genes in the absence of azoles (Supplementary Figures 4A and 4B). While in the presence of azoles, we clearly see an upregulation of *cyp51 genes* based on drug response, no upregulation was seen due to genotype (Two-Way ANOVA, interaction p-value >0.5, Supplementary Figures 4A and 4B). We also did not see a change in mRNA levels of transcription factors, *srbA* and *atrR*, that are mainly associated with *cyp51a/b* expression. (Supplementary Figure 4C and 4D).

### Loss of *afu-182* results in an azole recalcitrant infection in a murine model of invasive pulmonary aspergillosis

To test the role of *afu-182* during infection, we used a corticosteroid model of invasive pulmonary aspergillosis (39). We challenged mice with 2 x 10^6^ spores of the respective strains and monitored mice for 14 days for survival (Figure 4A). We did not see a difference in survival rates across strains (Figure 4B, log-rank test, p-value 0.1166 between WT andΔ*afu-182* strains). Also, we did not see a qualitative difference in fungal growth or histopathology as determined by GMS and H&E staining, respectively (Supplementary Figure 5). We next assessed the impact of *afu-182* loss on azole treatment outcome. Posaconazole was selected due to its higher bioavailability in the absence of grapefruit juice compared to voriconazole in mice (40, 41). Mice were challenged as previously described (39) and were treated with 5mg/kg of posaconazole at 24, 48 and 72 hours of infection (Figure 4C) . Mice challenged with Δ*afu-182* strains had significantly more adverse outcomes compared to mice infected with the WT strain (p = 0.0103, log-rank test) (Figure 4D). Thus, the Δ*afu-182* strain that is as virulent as WT in the absence of drug is recalcitrant to treatment in the presence of the azole drug. Additionally, we challenged and treated mice and harvested lungs after 72 hours (Figure 4E, schematic). We measured the fungal DNA as previously described (40). Our data indicate a significantly higher fungal burden in mice challenged with the Δ*afu-182* strain (p=0.0159, Mann-Whitney U-test) and a significantly reduced fungal burden in mice challenged with OE-182 strain (p=0.0159, Mann-Whitney U-test) upon azole treatment (Figure 4F); however, survival of animals infected with OE-182 strain and treated with posaconazole was not different compared to WT (p=0.4065, log-rank test, Figure 4D).

**Figure 4.**
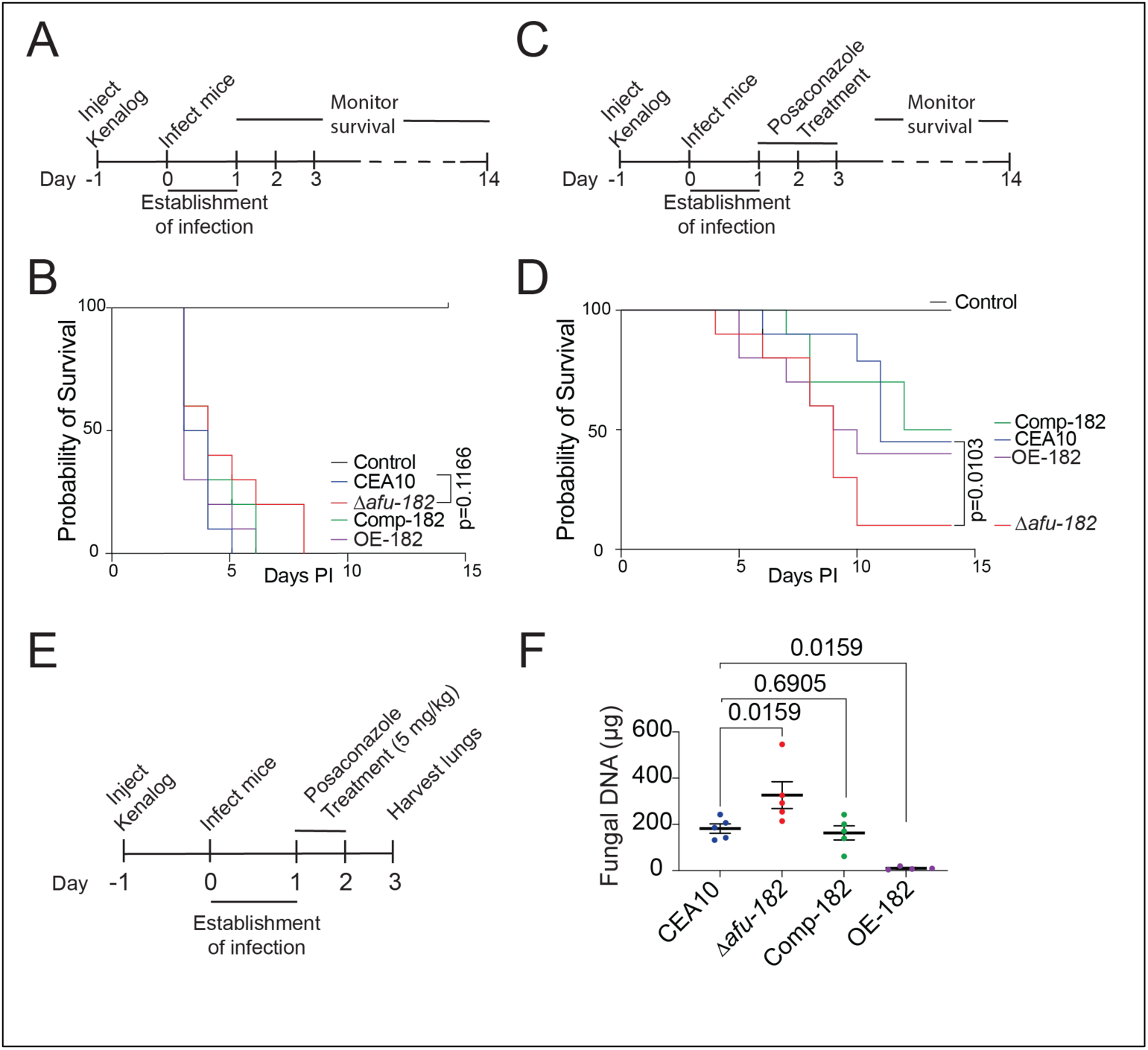
*afu-182* plays a role in azole treatment outcomes but is indispensable for virulence in a corticosteroid murine model of IPA. (A) Schematic showing experimental details for animal infection. (B) Kaplan-Meier plot of animal survival probability for animals infected as in A. No significant difference was observed (p=0.1166 WT vs. Δ*afu-182*, log-rank test). (C) Schematic showing experimental details for animal infection and azole treatment. (D) Kaplan-Meier plot of animal survival probability for animals infected as in C. A significantly higher mortality (90% vs 55%) was observed in animals infected with the Δ*afu-182* strain compared to animals infected with the WT strain (p=0.0103, log-rank test) upon azole treatment. No difference was observed in the mice infected with the OE-182 strain compared to WT (40% mortality vs 50% mortality, p=0.4065). (E) Schematic showing experimental details for infection and treatment. (F) Mice were infected and treated as in F. Fungal DNA was quantified as described in Methods. Significantly more fungal DNA was recovered from animals infected with the Δ*afu-182* strains (p = 0.0159, Mann-Whitney U-test), and significantly less DNA was recovered from animals infected with the OE-182 strain (p = 0.0159, Mann-Whitney U-test) after posaconazole treatment. No difference was observed in fungal DNA isolated from animals infected with WT or comp *afu*-*182* strains (p=0.6905).

### Sub-MIC azole exposure results in increased azole Minimum inhibitory concentration in an *afu-182* dependent manner

Lower drug concentrations at the infectious lesions compared to serum drug level is a challenging problem in treating fungal infections and fungus might be exposed to sub-MIC drug concentrations during infections. To determine the role of *afu-182* mediated sub-MIC response in the development of drug resistance i.e., change in MIC, we exposed WT and Δ*afu-182* strains to low dose azole at 0.1 μg/ml voriconazole every 48 hours for 12 generations in ten independent lineages (Supplementary Figure 6). After 12 generations, we see an increase in voriconazole MIC only in Δ*afu-182* strain in 6 lineages out of 10, but never in the WT strain, indicating that sub-MIC azole adaptation leads to an increase in MIC as measured by broth microdilution or E-Test strip in an *afu-182* dependent manner (Figures 5A and 5B). To further see the growth difference on solid medium we grew the adapted strains at varying spore densities (10^5^-10^2^) in presence of 0.5μg/ml voriconazole and see robust growth in Δ*afu-182* adapted strains but not in WT adapted strain (Figure 5C, red arrows). We did observe increase in sub-MIC (0.25μg/ml) growth of WT adapted strain (Figure 5C, orange arrow) but no discernable growth was observed at 0.5μg/ml voriconazole. We further tested the levels of azole target genes *cyp51a* and *cyp51b* and did not see a difference in RNA levels, indicating that the change in resistance in voriconazole adapted strain is not associated with transcriptional regulation of azole target genes (Supplementary Figure 7).

**Figure 5.**
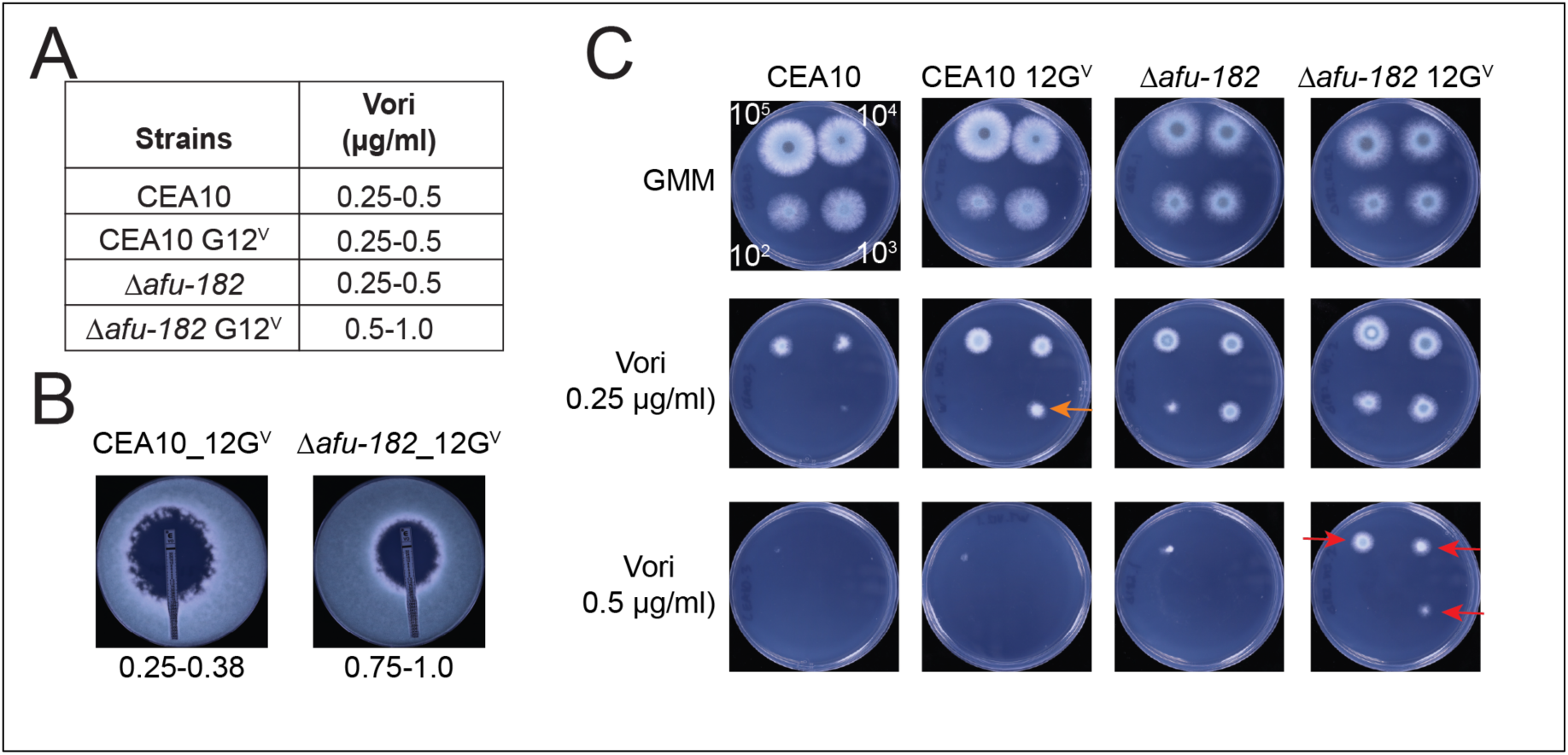
Continuous exposure to low level azoles leads to azole drug resistance in an afu-182 dependent manner. (A) Broth microdilution was used to determine the voriconazole MIC for the un-adapted and adapted strains (G12^V^ – 12 defines the generation and V defines adaptation in presence of voriconazole). Only the Δ*afu-182* strain exposed to low level voriconazole show increase in MIC. (B) E-test strip was used to assess the MIC of voriconazole of the adapted strains. Like broth microdilution, E-Test showed increase in MIC in Δ*afu-182* adapted strains. MIC values represented at the bottom as μg/ml. (C) Serially diluted spores (10^5^-10^2^) were spot inoculated on GMM or GMM + voriconazole (0.25μg/ml) or (0.5μg/ml) and grown for 48 hours at 37°C. The WT adapted strain showed increased fungal growth in the presence of the sub-MIC concentration of voriconazole (orange arrow), whereas the Δ*afu-182* adapted strain showed increased fungal growth at 0.5μg/ml (red arrows).

### *afu-182* regulates the RTA family of proteins in the presence of azoles

To determine the effects of azole on global transcription in WT on solid medium, we compared the transcriptomic profile of WT strain in the presence and absence of azoles with a stringent threshold change of 4-fold or more. We identified 4 genes (*afub_000050*, *afub_000060*, *afub_049700* and *afub_046760*) that are upregulated 4 fold or more in presence of azole in WT (Supplementary Figure 8) that contain a domain with predicted functions in stress response and belong to the RTA1family of proteins (32). In *Saccharomyces cerevisiae*, members of this family are involved in resistance to 7-<u>a</u>minocholestrol (RTA), resistance to <u>m</u>olasses (RTM), and lipid and heme transport (42). Previous report has suggested that a member of the RTA1 family is the most increased transcript in response to itraconazole in clinical samples (43). In *A*. *fumigatus*, we identified 23 genes in the genome encoding RTA1 family proteins. Interestingly, most of these genes are expressed at very low RNA levels (FPKM <5 for 17 genes in the glucose minimal medium conditions examined) and thus did not make the cut-off for global transcriptomic analysis. We selected 12 genes that showed significant change (2-fold or more) in transcript level and have an FPKM value >1 in GMM samples and performed qPCR. Comparative analysis revealed that all 12 gene transcripts were increased in the presence of azoles in WT (Figure 6).

**Figure 6.**
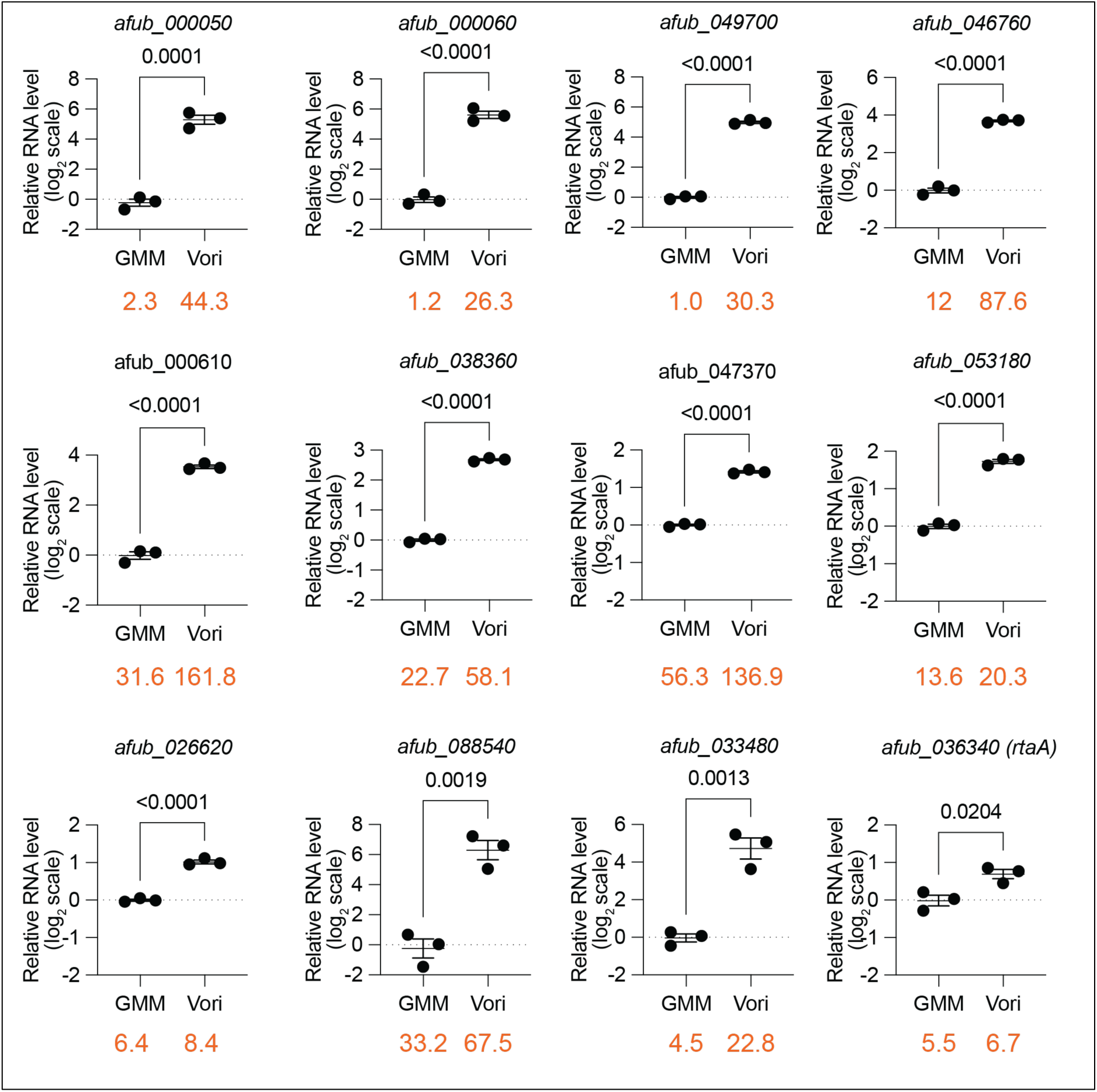
Genes encoding RTA1 domain proteins are upregulated in response to voriconazole. Quantitative reverse transcription PCR was used to determine the mRNA levels of RTA1 genes in the presence and absence of azoles. Out of 23, we selected 12 genes that have an FPKM value >1. Eleven of the 12 genes showed significant upregulation in the presence of azoles (p<0.05, two-tailed Student’s t-test). Importantly, *rtaA,* which plays a role in polyene response but not azole response, showed the lowest upregulation (1.6-fold, p=0.0204, two-tailed student’s t-test). Numbers in orange represent the FPKM values from the RNAseq data set.

To determine the role of *afu-182* in regulating fungal azole response, we compared the transcriptome of WT and Δ*afu-182* strain and saw 15 genes to be upregulated and 10 genes to be downregulated in Δ*afu-182* strain by 4-fold strain compared to WT in the presence of azoles out of about 10,000 variables (Figure 7A). Interestingly, two genes of the RTA1 family were the most upregulated genes in the Δ*afu-182* strain compared to WT indicating a negative regulation (Figure 7A, orange arrows) were upregulated in presence of azoles in WT. In the absence of azoles, their expression was not different (Figure 7B).

**Figure 7.**
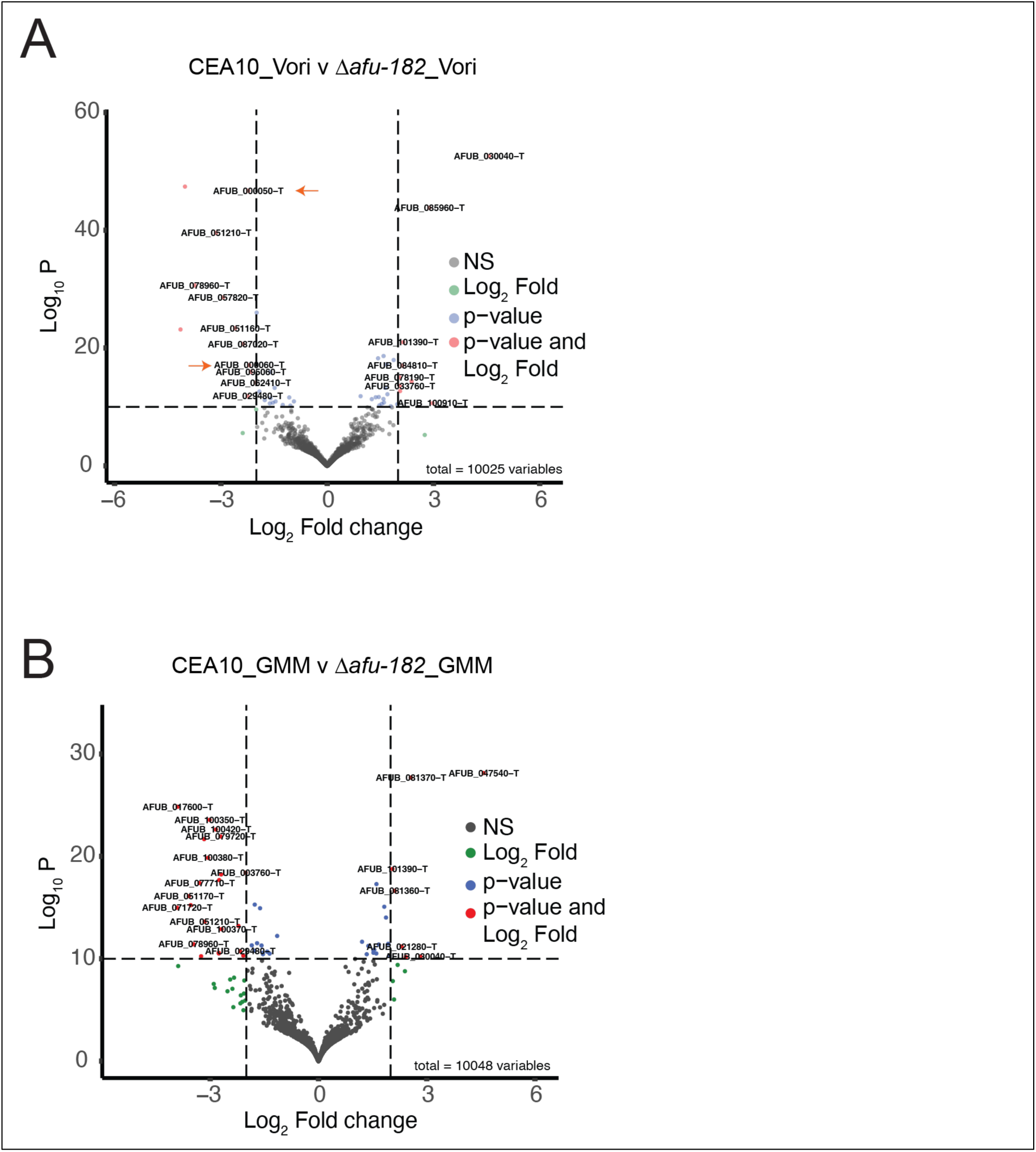
*afu-182* negatively regulates genes encoding for RTM1 proteins. Volcano plots showing the 2-log_2_ fold change (4-fold upregulated or downregulated). (A) Comparison between CEA10 azole and Δ*afu-182* azole. Global transcriptomic analysis revealed genes encoding RTM1 domain proteins *rtmA* (*afub_000050*) and *rtmB* (*afub_000060*) are significantly upregulated (orange arrows). (B) Comparison between CEA10 and Δ*afu-182* in GMM. No significant change in *rtmA* and *rtmB* level was observed. Quantitative reverse transcriptase PCR was used to assess the mRNA levels of (C) *rtmA* and (D) *rtmB*. Significant upregulation of *rtmA* (3-fold compared to WT, p=0.0005, Two-Way ANOVA followed by Tukey’s post-hoc analysis) and *rtmB* (4.6-fold compared to WT, p=0.0015, Two-Way ANOVA followed by Tukey’s post-hoc analysis) was observed in Δ*afu-182* strains compared to WT in the presence of voriconazole. No significant difference in *rtmA* and *rtmB* expression was observed in the absence of azoles.

Of the 23 genes in the RTA1 class, *afub_00050* and *afub_00060* are annotated as RTM1 proteins, a homolog of the *S. cerevisiae* gene providing resistance to molasses (44). We termed them as *rtmA* and *rtmB,* respectively.

### *rtmA* and *rtmB* regulate sub-MIC fungal azole response

To ascertain the role of *rtmA* and *rtmB* in fungal azole response, we deleted them in the CEA10 background. Additionally, we made a double null mutant Δ*rtmA*Δ*rtmB* strain. We also ectopically complemented *rtmA* and *rtmB* single mutants. Both the Δ*rtmA* (31% decrease in growth ratio, p=0.0012 One-Way ANOVA) and Δ*rtmB* (42% decrease in growth ratio, p<0.0001, One-Way ANOVA) strain showed reduced fungal growth in response to voriconazole (Figures 8A-B, and 8C-D), whereas the Δ*rtmA*Δ*rtmB* strain also showed significant growth attenuation in the presence of voriconazole (Figures 8E and 8F, 46% decrease in growth ratio p<0.001, One-Way ANOVA), indicating a combined role of these genes in regulating fungal azole response, where *rtmB* is the major driver of the azole response phenotype. The complement strains reverted to the WT levels, indicating a causal role of these genes in fungal azole response downstream of *afu-182* (Figures 8A-D, inset – macro lens images). Further studies are needed to determine the functional redundancy of other genes in this family; however, recently, an RTA1 family protein, *rtaA,* was shown to play no role in azole response (45). The functions of these genes appear to be diverse.

**Figure 8.**
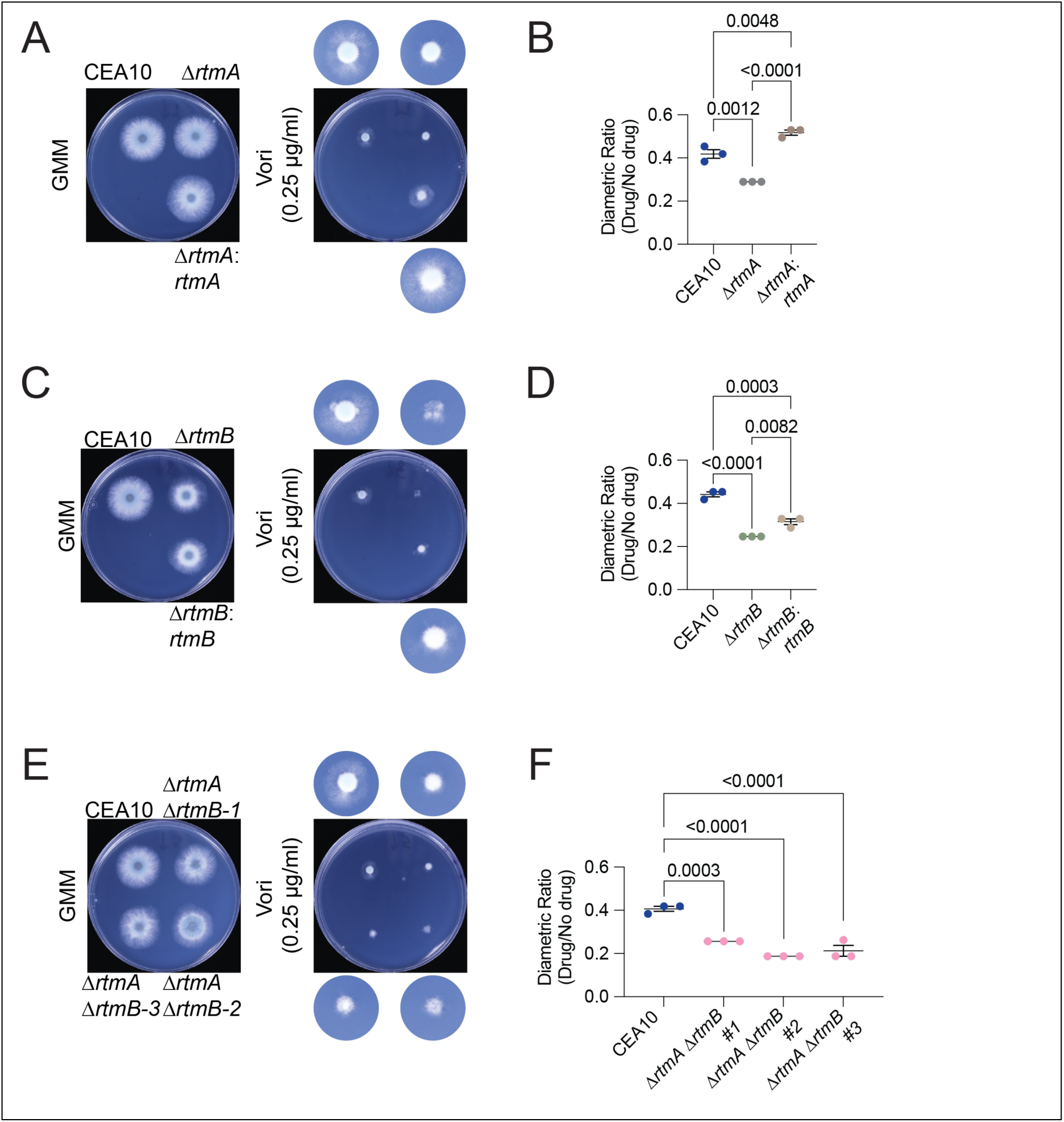
*rtmA* and *rtmB* play a role in fungal azole response. AF strains (A) Δ*rtmA* and comp-*rtmA*, (C) Δ*rtmB* and comp-*rtmB* and (E) Δ*rtmA* Δ*rtmB* were point inoculated in the absence or presence of 0.25 μg/ml of voriconazole. The plates were incubated at 37°C for 48 hours and photographed. The inset shows images taken with a macro lens. After 48 hours, colony diameter was measured and growth ratio of drug over no drug for (B) Δ*rtmA* and comp-*rtmA*, (E) Δ*rtmB* and comp-*rtmB* and (H) Δ*rtmA*Δ*rtmB* (three independent deletion strains) were plotted. One-way ANOVA was used to assess the difference in the mean. Numerical p-values are represented based on Tukey’s post-hoc analysis to compare the means of all groups.

## Discussion

While increasing azole resistance in *A. fumigatus* is a major concern, azole-susceptible strains (ASAF) still cause most infections (46–48). Agricultural use of azoles is a major source of *cyp51* mediated environmental azole resistance (49). This hints at a broader response to AF drugs, especially in ASAF, during infections. Here, we show that the long non-coding RNA *afu-182* negatively regulates fungal response to sub-MIC azole drug levels in ASAF (Figure 2), and deletion of *afu-182* is not associated with a change in MIC (Supplementary Figure 2D). *afu-182* was previously identified as an anti-sense ncRNA to *insA* (32) that was recently shown to play a role in azole drug resistance (27); however, the role of *afu-182* has been obscure. (32, 33). Here, we show that *afu-182* lncRNA regulate fungal response to sub-MIC azole drugs.

Fungal response to drugs has been characterized based on changes in genetic sequence (i.e., resistance) or non-genetic mechanisms (tolerance, persistence, and hetero-resistance) (28, 30, 31). We see robust growth of Δ*afu-182* strain at azole concentrations approaching MIC but all strains fail to germinate at 2x MIC concentration (Supplementary Figure 9). Thus, the Δ*afu-182-*mediated phenotype at sub-MIC azole concentration is likely a stress adaptation mechanism to tolerate azole drugs. Furthermore, the Δ*afu-182-*mediated sub-MIC growth phenotype is independent of the strain’s background and is conserved in related species *A*. *nidulans* (Supplementary Figure 3).

We also showed the increased fungal growth upon azole treatment in fungal surface attached cultures, thus indicating effects are independent of spore germination (Figure 2F). Drug resistance is a classical feature of the biofilms (50), and even though we saw increased attached fungal biomass for the Δ*afu-182* strain upon azole treatment (Figure 2F), it is possible that different mechanisms are playing roles in this distinct growth conditions. For example, metabolic adaptation, drug binding to the ECM, and oxygen gradients have all been shown to play a role in biofilm mediated drug resistance (51, 52). Additionally, biofilms can lead to persister cells (53). Further research will uncover if distinct mechanisms confer difference in azole response in surface attached cultures.

Different AF isolates show differential *afu-182* levels, implying that *afu-182* might be involved in fungal adaptation to various stresses, though in our murine model, it was not required for virulence (Figure 4B and Supplementary Figure 5). Thus, the evolutionary significance of *afu-182* and its role in adaptation against other stresses in AF is currently under investigation. Importantly, the Δ*afu-182* strain shows increased fungal growth only in response to azole stress and does not play a role in response to the polyene and echinocandin classes of drugs (Supplementary Figure 10). Over 90% of infections are still caused by ASAF, with treatment failure rates approaching 50-90% (54, 55). Thus, it is imperative to better understand the fungal response to azole drugs.

To understand the role of *afu-182* in fungal azole response *in vivo*, we used a steroid model of IPA. In the absence of azole drugs, the Δ*afu-182* strain is pathogenic and equally virulent as WT (Figure 4B); however, in the presence of 5mg/kg of posaconazole, our data show increased virulence in the Δ*afu-182* strain as measured by survival analysis and fungal burden analysis (Figure 4D and 4F). The overexpression strain did not show a change in survival rate compared to WT when treated with azole drugs (Figure 4D); however, it showed a 90% reduction in fungal burden, highlighting the role of *afu-182* in azole treatment outcomes (Figure 4F). In a steroid model of invasive aspergillosis, deaths due to exacerbated immune response have been reported (55), and it is plausible that this is the case in OE-182*-*infected azole-treated animals. In vitro, we see that *afu-182* regulates sub-MIC response to all azoles, including posaconazole (Figures 2A and 2D). Like *in vitro* response, it is possible that low levels of azole exposure in fungal lesions lower the *afu-182* RNA levels that allow the AF to grow better in the presence of the drug, thus increasing fungal burden in Δ*afu-182* strain and lower burden in OE-182 strain. As drug resistant isolates are recovered from patients during azole treatment if IA or IPA (56), this sub-MIC azole adaptation may eventually lead to drug resistance. To test this *in vitro*, we adapted the WT and the Δ*afu-182* strain in presence of 0.1μg/ml of voriconazole (Supplementary Figure 6) and observed a 2-fold change in MIC of voriconazole only in Δ*afu-182* strain but not in WT strain (Figures 5A and 5B). Thus, it is possible that during the long-term infection low dose interactions leads to *afu-182* mediated fungal adaptation, and this adaptation eventually leads to increase in MIC above clinical breakpoints (Figure 9).

**Figure 9.**
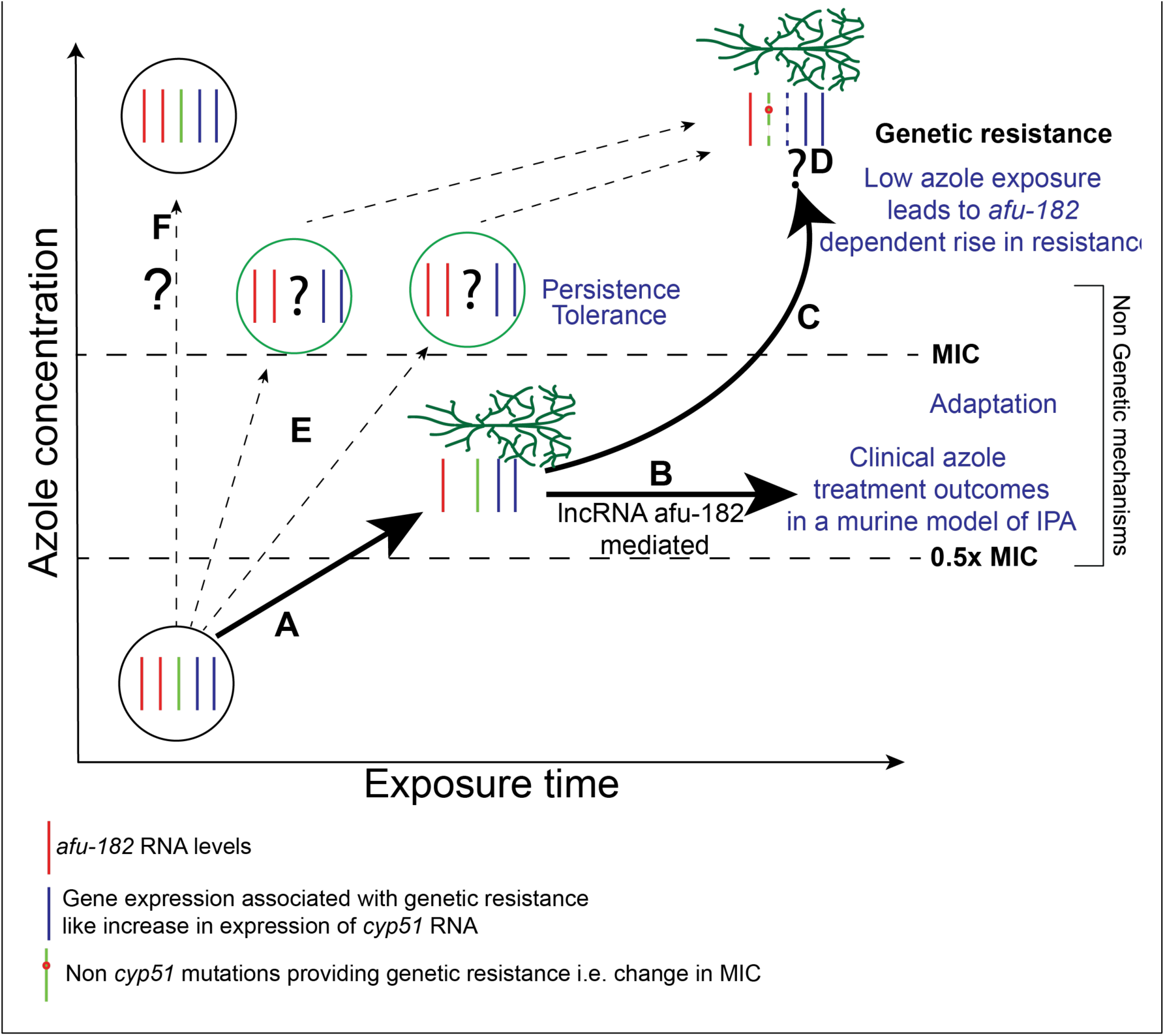
AF responds to sub-MIC azole drugs by regulating *afu-182* levels resulting in azole drugs resistance. (A) Sub-MIC azole exposure lowers the *afu-182* levels. (B) This adaptation affects the clinical outcomes in an animal model of IPA treated with azole drugs. (C) Continuous exposure to low level drugs that is possible during infection or environmental use leads to increase in MIC. (D) The mechanistic pathway controlling *afu-182* mediated resistance are unknown and under investigation; however, appear to be independent of Cyp51 transcriptional regulation. (E) Acute exposure of supra-MIC drugs is linked to tolerance and persistence, and it is currently unknown if that plays a role in drug resistance or (F) Spontaneous mutations can lead to azole drug resistance.

Drug concentration at the site of infection is a major indicator of successful therapy (57). Crucially, the inherent rise of drug resistance in 43% of isolates could not be attributed to changes in the *cyp51A* sequence (56), indicating non-*cyp51* mechanisms at play. It is known that continuous exposure to low level of drug leads to resistance whereas transient exposure to high concentration of drugs lead to tolerance (58, 59). How fungus adapts to anti-fungal stress before genetic changes associated with resistance occur is still unknown. Many studies have shown differences in serum level vs tissue level concentrations of anti-fungal drugs (57). Posaconazole has been shown to accumulate in lipid rich membrane of alveolar cells in high concentration in healthy subjects (60); however, the lesion concentration might still be low. Also, during infection tissue necrosis may limit interaction of fungal cells and posaconazole rich cell membrane thus limiting lesion concentrations (61). Thus, drug treatment challenges may lead to low dose exposure in the fungal lesions in patients with IPA even when serum concentrations are high, leading to lower *afu-182* RNA levels, stress adaptation and rise in inherent drug resistance in AF during infection (Figure 9).

Different isolates of *A. fumigatus* have been previously shown to differ in virulence (62). Some of the factors attributed to this difference are nutrient plasticity and response to light and oxygen (62–64); however, how different azole-susceptible isolates respond to azole drugs and their virulence potential in relation to azole treatment requires further research. *afu-182* regulates fungal azole response in ASAF irrespective of the strain’s background, as indicated by deletion of *afu-182* in the laboratory isolates CEA10 and Af293 backgrounds (Figures 2 and 3D-E) (37). Additionally, deletion of *afu-182* in the environmental isolate 47-57 also leads to differential fungal response to sub-MIC concentrations of azoles (Figures 3D and 3F).

The emergence of new strains provides a unique challenge in studying *Aspergillus* pathogenicity, but *afu-182* appears to play a role irrespective of the strain’s background. One possible reason for this might be the mechanisms by which ncRNA functions. LncRNA provide stress adaptation in the absence of evolution that is associated with genomic changes (65). A rapid adaptation will increase the likelihood of fungal survival during stress, leading to DNA changes or the accumulation of beneficial mutations. A recent study identified 1089 novel long non-coding RNAs; eleven of which negatively regulate fungal response to azole drugs; however, *afu-182* fell below the size threshold identified in this study (66). How *afu-182* regulates fungal response is not yet clear but it is independent of *cyp51* regulation at the transcriptional level (Supplementary Figures 4A and 4B). Also, mRNA levels of *srbA* and *atrR*, the TFs that regulate *cyp51* did not change in Δ*afu-182* strain (Supplementary Figures 4C and 4D) indicating either the effect in post-transcriptional, downstream of these TFs or completely independent of these known mechanisms.

As long non-coding RNAs function in diverse ways, we used a transcriptomic approach to identify the genes directly or indirectly regulated by *afu-182* at the RNA levels. In the presence of azoles, we see the RTA1 family of proteins upregulated in WT (Figure 6). This is a 7-transmembrane domain family of proteins and has been associated with lipid and heme export and resistance to 7-amonicholesterol in *S. cerevisiae* and *Cryptococcus neoformans* (42, 67) and consists of 23 members in *A. fumigatus*. Recently, *rtaA* (Afu3g12830, CEA10 homolog - *afub_036340*) was shown to play a role in response to the polyene class of drug, amphotericin B and not azoles (45). Here, we identified *rtmA* and *rtmB,* two divergent genes downstream of *afu-182*, situated next to each other to regulate fungal azole response (Figure 8). Both *rtmA* and *rtmB* are negatively regulated by *afu-182* (Figures 7); however, *rtmB* mutations might be the primary contributor to the azole response phenotype, as the double deletion strains show nearly the same inhibition as Δ*rtmB* (Figures 8D and 8F). As both these genes are next to each other, it is not clear whether *afu-182*, by itself or in coordination with other complex members, if any, regulates the promoter region of these genes. It is also possible that these genes evolved due to a duplication event to maintain tight regulation for stress adaptation. If *afu-182* regulates fungal azole response via direct regulation of RTM/RTA1 domain-containing proteins is an active and ongoing area of research.

Importantly, *rtmA* and *rtmB* belong to the accessory genome of AF and were detected in only 7 of the 300 AF genomes previously analyzed (68). Thus it is possible that the phenotypes observed in Δ*rtmA* and Δ*rtmB* are correlative with Δ*afu-182* phenotypes as these genes are absent in Af293 strains of AF (68) but we still see the Δ*afu-182* mediated sub-MIC phenotype in Af293 background (Figures 3D and 3E). It is also possible that in absence of *rtmA* and *rtmB* genes, other RTA1 family proteins play a similar role in Af293. More research will uncover the function of this family of proteins in AF in the future.

Long non-coding RNAs mainly function through their secondary structure and are difficult to predict based on the nucleotide sequence alone (69). Here, using synteny, we identified *afu-182* homolog in related fungal species *A. nidulans*. Deletion of *afu-182* homolog, *an-182,* also lead to differential sub-MIC azole response in *A. nidulans,* showing its role in multiple Aspergilli (Supplementary Figure 3). Whether *afu-182* homologs play a role in the phytopathogens *A*. *flavus* and *A*. *parasiticus* is unknown and needs more research; complex approaches are needed to identify the homologs of *afu-182* in other fungal pathogens. However, our data clearly show that long non-coding RNA, *afu-182,* regulates fungal adaptation in presence of sub-MIC azole concentration in AF and AN irrespective of the strain’s background.

In conclusion, the lncRNA *afu-182* regulates fungal response to azole drugs in both ASAF and the related fungal species AN. This adaptation is beneficial for AF to gain azole resistance. This effect, led to identification of *rtmA* and *rtmB* genes in RTA1 family of protein that regulate fungal azole response and whose transcription is negatively regulated by *afu-182* in CEA10 strain. As MIC is a poor predictor of azole treatment outcomes in ASAF strains, further characterization of *afu-182* levels will help us determine the fungal response to azole drugs *in vivo* that might contribute to inherent rise in azole drug resistance during azole treatment. Given advancements in RNA biology, *afu-182* also presents a target to augment the efficacy of current azole drugs.

## Materials and Methods

### Fungal strains

*Aspergillus fumigatus* (AF) strain CEA10 or CEA17 was used as the host strain to generate subsequent strains. For *A. nidulans*, A1145 (FGSC) and GR5 (FGSC) were used as the host strains to generate subsequent strains. For *A*. *nidulans*, the strains were supplemented with Uridine (1.26g/L), Uracil (0.56g/L), pyridoxine (0.05μg/ml), p-amino-benzoic acid (PABA) (5μM) and riboflavin (0.1μg/ml). All experiments with stationary fungal growth were performed at 37°C in the presence of 5% CO_2_. Liquid biomass and germination assay were performed in the absence of supplied CO_2_. All strains were stored as 30% glycerol stocks in -70°C for long-term storage. The list of strains used and generated is listed in Supplementary Table 1. All primers and crRNAs used in this study are listed in supplementary tables 2 and 3 respectively.

### Δ*insA* strain construction

The *insA* deletion strain was created using homologous recombination. Fusion PCR was used to create the DNA deletion cassette. Briefly, the 5’ and 3’ fragments of *insA* were amplified with primers RAC 815 and RAC 816, and RAC819 and RAC820, respectively. A gene encoding for *pyrG* was amplified from pJW24 (70) using primers RAC817 and RAC818. The three fragments were fused together using primers RAC 821 and RAC 822. The resulting DNA fragment was transformed into the CEA17 strain via PPEG-mediated transformation of protoplast (71). Strains were checked with PCR, Southern blot for single integration and qPCR for lack of detectable RNA. This strain does express *afu-182* at the native locus.

### Complementation plasmid construction

A plasmid encoding for *pyrithiamine* resistance, BS309, was created as follows. Two oligo fragments (SD37 and SD38) encoding for 5’ SacI overhang-AscI-BglII-MluI-Pac1-SacII-3’ overhang were ordered from IDT and were allowed to anneal in the thermocycler. The resulting dsDNA fragment with SacI-SacII overhangs was ligated into pBS KS (+) plasmid previously digested with SacI and SacII, resulting in plasmid BS309 intermediate. A DNA fragment encoding for pyrithiamine resistance was amplified from pSD51 with primers RAC3280 and RAC3154 expressing an XhoI restriction site (72). The DNA fragment was digested with XhoI and ligated into BS309 intermediate previously digested with SalI, resulting in the plasmid BS309.

### Plasmid construction for Comp-*insA* with cis acting *afu-182*

A DNA fragment consisting of ∼1kb 5’ region of *insA* along with *insA* and 485bp after the transcription end site (as determined by RACE) expressing AscI and PacI restriction sites at the 5’ and 3’ ends, respectively, was amplified with primers RAC3589 and RAC3591 using Takara Prime Star HS polymerase per manufacturer’s recommendation. The resulting DNA fragment was digested with AscI-Pac1 and was ligated into BS309, resulting in comp-cis *insA* plasmid.

### Plasmid Construction for Comp-*insA* without cis-acting *afu-182*

A DNA fragment consisting of 5’ region of *insA* along with *insA* and 74bp after the transcription end site (without ncRNA *afu-182*) expressing AscI and PacI restriction sites at 5’ and 3’ ends respectively was amplified with primers RAC3589 and RAC3590 using Takara Prime Star HS polymerase per manufacturer’s recommendation. The resulting DNA fragment was digested with AscI-Pac1 and was ligated into BS309, resulting in comp-without cis-182 *insA* plasmid.

The resulting plasmids were transformed into the Δ*insA* strain using PEG-mediated transformation of protoplast as previously described (71).

### Δ*afu-182* strain construction

*afu-182* was deleted from the strain CEA17 (*pyrG*-) using CRISPR-Cas9 as previously described (73). A homology-directed repair cassette was amplified from plasmid pSD38.1 using primers SD149_182 homology F and SD150_182 homology. Briefly, 5μl of 100mM crRNAs (on either side of *afu-182*) were combined with 5μl of 100mM tracrRNA and 5 μl of nuclease free duplex buffer (IDT), heated to 95°C for 5 min and allowed to cool to room temperature, yielding 33mM gRNA complex. Then, 1.5 μl of each guide RNA complex on either side of *afu-182* were combined with 1.5μl of Cas9 (1mg/ml) and 22μl of Cas9 working buffer as previously described (73). The samples were allowed to combine at RT for 5 min to allow the assembly of the RNP complex. RNP complex and repair cassette were transformed via PEG-mediated transformation of the CEA17 strain as previously described (71, 73). The same gRNAs and HDR were also used to delete *afu-182* in the Af293.1 strain and 47-7 strain (62). The sequence of crRNAs and primers used is provided in Tables S2 and S3.

### Comp-182 strain construction

*afu-182* was put back at its genomic location using CRISPR-Cas9. Briefly, the *afu-182* fragment was amplified with primers SD76 and 164 using CEA10 gDNA as template. A gene encoding for pyrithiamine resistance was amplified from plasmid pSD51.1 (72) using primers SD23 and SD165. Both fragments were fused using primers SD183 and SD166, generating the HDR cassette. A single gRNA (gRNA 13 SD 1) targeting the linker sequence in the *pyrG* gene was used for double-strand break, and HDR was used to insert *afu-182* back at the locus.

### OE *afu-182*pyrG plasmid construction

The *A*. *nidulans gpdA* promoter was amplified from strain FGSC A4 (FGSC) using primers SD134 and SD103. The *afu-182* fragment was amplified from CEA10 gDNA using primers SD104 and SD105. Subsequently, these fragments were fused using primers SD135_EcoRI and SD139_NotI and digested with EcoRI and NotI. The resulting fragment ligated into plasmid pSD21 previously digested with EcoR1 and Not1, yielding plasmid OE*afu-182*pyrG. The *afu-182* RNA expression starts at the +1 transcription start site (TSS).

### CEA10 OE *afu-182* strain construction

CRISPR-Cas9 was used to target the OE *afu-182 pyrG* HDR cassette to the safe-haven *atf4* locus (74) as previously described (64). The HDR was amplified from plasmid OE *afu-182pyrG* using primers SD162_OE_homology-F and SD163_OE_homology-R. gRNA was assembled as described above for the Δ*afu-182* strain. A single guide (atf4_gRNA7) was used to initiate a double-stranded break targeting the *atf4* locus (74).

Polyethylene glycol-mediated fungal transformation was done as described above. All resulting transformants were checked with PCR and confirmed with Southern blot and qPCR for gene expression.

### *Aspergillus nidulans* Δ182 (Δan-182) construction

A DNA sequence homologous to *afu-182* was identified 240bp after the translational stop site of the *insA* homolog gene AN4465 (32). The resulting 200bp sequence was deleted from the *A. nidulans* genome using CRISPR-Cas9 in the A1145 strain using the *pyrG* marker or the GR5 strain using the *pyroA* marker as previously described (73). The HDR cassette for A1145 was amplified using primers SD181 and SD182 that amplified the sequence of orotidine 5’-phosphate decarboxylase (75) gene from *A*. *parasiticus* using plasmid pSD38.1 (76) as template. The HDR cassette for GR5 was amplified with SD 352 and SD 353, which amplified the pyridoxine biosynthesis gene (77) from *A. fumigatus* using CEA10 as the DNA template.

### Δ*rtmA*, Δ*rtmB and ΔrtmAΔrtmB* strains construction

Δ*rtmA* and Δ*rtmB* strains were constructed using the CRISPR-Cas9 protocol as previously described for the Δ*afu-182 strain* (64). The HDR cassette primers and crRNA sequences are provided in Table S1.

Δ*rtmA* and Δ*rtmB* strains were constructed in the CEA10 background using the pyrithiamine resistance marker, *ptrA* from pSD51.1. For Δ*rtmA*, crRNA 50L and 50R were used, for Δ*rtmB*, crRNA 60L and 60R were used. For double deletion strains, crRNAs 60R and 50R were used. HDRs were amplified using primers SD151 and SD152 for Δ*rtmA* deletion; SD153 and SD154 for Δ*rtmB* deletion and primers SD152 and SD153 for double deletion.

### Complement *rtmA* and complement *rtmB* strain construction

Hygromycin resistance gene expressing plasmid: Hyg resistance gene was amplified using the primers RAC2939 and RAC2940 from the plasmid pBC Hygro (FGSC) and was ligated into pJET1.2 (Thermo Fisher). The fragment was digested using XhoI-XbaI and was ligated into the plasmid BS309 (see above), previously digested with XhoI and XbaI, resulting in plasmid BS311.

### Plasmid construction for Comp *rtmA* strain

The *rtmA* fragment, along with the promoter and terminator region, was amplified (CEA10 gDNA template) using the primers SD738 and SD739 with Asc1 and Pac1 restriction sites, respectively, using NEB Q5 hot start polymerase per manufacturer’s recommendation. The resulting DNA fragment was digested with AscI and PacI and was ligated into the plasmid BS311 previously digested with AscI and PacI, resulting in the Comp-*rtmA* plasmid.

### Plasmid construction for comp-*rtmB* strain

The *rtmB* fragment, along with the promoter and terminator regions, was amplified (CEA10 gDNA template) using the primers SD740 and SD741 with Asc1 and Pac1 sites, respectively, using NEB Q5 hot start polymerase per manufacturer’s recommendation. The resulting DNA fragment was digested with AscI and PacI and was ligated into BS311 previously digested with AscI and PacI, resulting in the Comp-*rtmB* plasmid.

Comp-*rtmA* and comp-*rtmB* plasmids were transformed into Δ*rtmA* and Δ*rtmB* strains, respectively, as described above.

### Spot assays

Plates containing solid GMM were spot inoculated with serially diluted AF spores (10^5^-10^2^ spores) and grown at 37°C as indicated. Anti-fungal drugs (azoles, amphotericinB and Caspofungin) were used at the indicated concentrations. The colony diameter was measured and reported as the growth ratio, dividing the colony diameter in the presence of azole by the colony radial diameter in GMM. All experiments were done at least in triplicates and were repeated three times. Statistical analyses (t-test or one-way ANOVA) were done to determine the significant differences using GraphPad Prism.

### Biomass assay

Spores were grown in 50 ml liquid GMM (1 x 10^6^ spores/ml) at 37°C for 24 hours in the presence or absence of 0.2 µg/ml voriconazole with continuous shaking at 250rpm. Mycelium collected after 24 hours was lyophilized, and the biomass was weighed. Statistical difference between means was assessed with One-way ANOVA followed by Tukey’s post hoc analysis to compare individual groups using GraphPad Prism.

### Biofilm formation assay

AF spore suspension (1.5 x 10^5^ spores/ml) in liquid GMM was inoculated in three biological replicates in a 12-well plate. The plates were centrifuged at 250g for 10 min to settle the conidia at the bottom of the wells and incubated at 37°C for 12 hours. Then, the wells were washed twice with PBS to remove planktonic cells. The biofilms at the bottom of the wells were further allowed to grow in the presence of 0.2 μg/ml Voriconazole for 8 hours before the crystal violet assay was performed as previously described (36). Absorbance was measured using a plate reader (Multiscan Sky, Thermo scientific) at 600 nm. All experiments were done in triplicate, and three independent experiments were performed. Statistical significance was determined with One-way ANOVA using GraphPad Prism.

### E-test assay

Five milliliters of GMM top-agar (0.75%) containing AF spores (1x 10^5^/5 ml) were spread on GMM plates and allowed to solidify. A voriconazole E-test strip (Biomerieux) was carefully laid in the center of the plate per the manufacturer’s recommendation. The plates were incubated at 37°C for 48 hours, and the minimum inhibitory concentration (MIC) was recorded. The experiment was done in triplicate for each strain and repeated three times.

### Broth Microdilution assay

AF spores collected from a 48-hour-old colony and (5 x 10^4^) were inoculated in liquid GMM in a 96-well plate containing voriconazole (16μg/ml – 0.015μg/ml) and incubated at 37°C for 48 hours. The visible growth of AF strains was observed in each of the wells, and the lowest concentration of azoles showing no growth was recorded as MIC. The experiment was repeated three times.

### Murine model of pulmonary aspergillosis, lung histopathology, and fungal burden

Outbred female CD-1 mice (Charles River Laboratories) with body weights of 18-21 g were housed (5 per cage) in micro-isolator cages. The mice were kept in a soft-lit, well-ventilated cubicle with a 12h:12h light/dark cycle. Mice were immunosuppressed with a single subcutaneous injection of 40 mg/kg body weight of Triamcinolone acetonide (Kenalog-10, Bristol-Myers Squibb) on day -1. Mice were briefly anesthetized with isoflurane and infected by intranasal instillation of 2 x 10^6^ spores in 40μL PBS. Ten mice were infected per AF strain for survival analysis, whereas five mice per strain were used for fungal burden analysis. The mice were observed for 14 days for their survival. For azole treatment, 5 mg/kg Posaconazole (Noxafil, oral suspension, Merck) diluted in PBS was used.

For fungal burden analyses, mice were treated with 5 mg/kg Posaconazole on day +1 and day +2, and their lungs were harvested on day +3. The mouse lungs were lyophilized overnight, homogenized with 2.3 zirconia beads, and lysed with LETS buffer (0.1M LiCl, 10mM EDTA pH 8.0, 10mM Tris-Cl pH 8.0, 0.5% SDS). Total DNA was extracted in an aqueous phase using phenol: chloroform: isoamyl alcohol and precipitated with isopropanol. Fungal burden was determined by calculating the levels of 18s rRNA encoding gene against a standard curve in 500ng total DNA using iTaq universal probe mix and hydrolysis probe as previously described (40). The statistical significance of survival analysis was determined using log-rank analysis, and fungal burden analysis was done using a non-parametric Mann-Whitney U Test.

For histological analysis, lungs were harvested from untreated mice on day +3 and were fixed in 10% formalin for 48 hours before microdissection at the Clemson Light Imaging Facility (CLIF) at Clemson University. Adjacent slides obtained from microdissection were stained with Grocott’s Methenamine Silver and Hematoxylin and Eosin stains as previously described (41). The stained slides were pictured with the Leica Laser Microdissection Microscopy System at CLIF.

### Voriconazole adaptation

Azole adaptation experiment was carried out for CEA10 and Δ*afu-182* strains by growing the strains in 0.1 µg/ml voriconazole for 12 generations at 37°C. Briefly, CEA10 and Δ*afu-182* strains were spot inoculated out of glycerol stock (0G) in glucose minimal media (GMM) plate containing 0.1 µg/ml voriconazole at 37°C for 48 hours. Spores (1G^V^) were collected in 0.01% Tween-80, and again spot inoculated onto fresh GMM plate containing 0.1 µg/ml voriconazole for 48 hours. The remaining 1G^V^ spores were stored as glycerol stock for further experiments in -70°C. This process was repeated for 12 generations.

### RNA extraction

For RNA sequencing, total RNA was extracted from AF strains CEA10, Δ*afu-182*, comp-182, and OE-182 strains. Briefly, 1 x 10^4^ spores were spot inoculated and incubated at 37°C for 48 hours on solid GMM plates in the absence or presence of voriconazole (0.15 µg/ml). The whole colonies from samples were excised with a clean scalpel and flash frozen in liquid N_2_. The frozen samples were homogenized with 2.3 mm zirconia beads in acid guanidinium thiocyanate phenol, and RNA was extracted using the acid guanidinium thiocyanate phenol chloroform (AGPC) method as described with some modifications (78). Briefly, the aqueous phase was mixed with ethanol to a final concentration of 35%. The resulting mix was passed through RNA-binding columns (Epoch Life Sciences), washed twice with 70% Ethanol and eluted in nuclease-free water. The RNA was treated with Turbo DNase (Thermo Fisher) per the manufacturer’s recommendation and purified using magnetic beads (Beckman Coulter). RNA quantity and quality were estimated using the AccuBlue Broad Range RNA Quantitation Kit (Biotium) and the Quantum Fluorometer (Promega), and via gel electrophoresis, respectively, before submission of the samples to LC Sciences.

Sample quality was checked by Agilent Technologies 2100 Bioanalyzer at LC Sciences. The poly(A) RNA sequencing library was prepared following Illumina’s TruSeq-stranded-mRNA sample preparation protocol. Poly(A) tail-containing mRNAs were purified using oligo-(dT) magnetic beads with two rounds of purification. After purification, poly(A) RNA was fragmented using a divalent cation buffer at elevated temperature. Quality control analysis and quantification of the sequencing library were performed using Agilent Technologies 2100 Bioanalyzer High Sensitivity DNA Chip. Paired-end sequencing was performed on Illumina’s NovaSeq 6000 sequencing system.

### RNA-seq analyses

Transcript expression was determined by first aligning RNA-Seq reads to the *A. fumigatus* A1163 genome obtained from FungiDB v68 (32) with the splice-aware aligner STAR v2.7.11a (79). The resulting SAM file for each experimental replicate was processed with samtools v1.19.2 (80) and featureCount v2.0.6 (81) to produce a table of read counts per annotated gene locus. The read counts were analyzed with DESeq v1.46.0 (82) and plotted with EnhancedVolcano v1.24.0 (83) in the R v4.4.2 statistical environment(84). The pipeline for analyzing these is archived in a GitHub repository (https://github.com/stajichlab/DhingraLab_ncRNA_Afumigatus) and Zenodo (85)

### Assessment of gene expression by RT-qPCR

For qPCR, 48-hour-old AF colonies grown at 37°C in solid GMM plates with or without voriconazole (0.15μg/ml) were extracted as described above. RNA samples were then treated with Turbo DNase per the manufacturer’s recommendations. Equal amounts of DNase-treated RNA were used for first-strand cDNA synthesis using MMLV reverse transcriptase (Promega) per the manufacturer’s recommendation. Quantitative PCR (CFX real-time PCR system - Bio-Rad) was done using the sso Advanced universal SYBR green supermix (Bio-Rad). Histone H4 was used as the reference gene for normalization. All primers used in the study are listed in Supplementary Table 2.

### Data Availability

The RNA-seq sequencing data and per-gene read counts are available from Sequence Read Archive associated to BioProject PRJNA1334547 and in Gene Expression Omnibus under accession GSE309441.

### Statistical analyses

GraphPad Prism 10 for macOS was used to perform all statistical analyses.

## Acknowledgements

We would like to acknowledge members of Dhingra laboratory and Eukaryotic Pathogen innovation center for discussions surrounding the experiments in the manuscript. The research in Dhingra lab is supported by start-up funds from the Department of Biological Science at Clemson University and National Institute of General Medical Sciences (NIGMS) award (P20GM146584) (PI - James Morris). We thank Dr. Robert Cramer at the Dartmouth Geisel School of Medicine (NIAID/NIH grants R01AI181215 and 2R0AI130128) for providing resources and critical reading of the manuscript. Analyses were performed on the UC Riverside High Performance Computing Cluster supported by grants from the National Science Foundation (DBI-1429826 & DBI-2215705) and the National Institutes of Health (NIH) (S10-OD016290). JES is a CIFAR fellow in the program Fungal Kingdom: Threats and Opportunities and supported by NIH grant R01AI130128.0. The Clemson University Genomics and Bioinformatics Facility is supported by the College of Science and Grants P20GM146584 and P20GM139769 Institutional Development Awards (IDeA) from the National Institute of General Medical Sciences of the National Institutes of Health. This work was supported, in part, by equipment housed in the Clemson Light Imaging Facility. We utilized a Leica Laser Microdissection system for brightfield imaging, and a Leica Pearl Automatic Tissue Processor and a Leica Histocore Biocut Rotary Microtome for histology preparation. Thank you to Mr. Avery Herren for assistance with histology. The Clemson Light Imaging Facility is supported by the Division of Research and EPIC COBRE P20GM146584. We thank Dr. Jarrod Fortwendel for the Fdel262 strain, Anne Strong for technical help and Dr. James Morris for the critical reading of the manuscript. All animal experiments were approved by Institutional Animal Care and Use Committee (IACUC approval # AUP2022-0236). Garret Kaufman was supported by MENTOR: Medical Enrichment Through Opportunities in Research Grant (1T35AI134643-01A1, PI – Dr. Kerry Smith). We acknowledge the support of veterinary and technical staff at the Godley-Snell Research Center for help with the animal studies. The funding sources had no role in the study design, data collection and interpretation, preparation of this manuscript or the decision to submit the manuscript.

## Notes

### Competing Interest Statement

The authors have declared no competing interest.

### Summary of Updates

Additional experiments added to show the role of afu-182 in fungal response to azole drugs.

## References

1. Morrissey CO, Kim HY, Duong T-MN, Moran E, Alastruey-Izquierdo A, Denning DW, Perfect JR, Nucci M, Chakrabarti A, Rickerts V, Chiller TM, Wahyuningsih R, Hamers RL, Cassini A, Gigante V, Sati H, Alffenaar J-W, Beardsley J. 2024. Aspergillus fumigatus—a systematic review to inform the World Health Organization priority list of fungal pathogens. Med Mycol 62:myad129.

2. Baddley JW. 2011. Clinical risk factors for invasive aspergillosis. Med Mycol 49 Suppl 1:S7–S12.

3. Latgé J-P, Chamilos G. 2019. Aspergillus fumigatus and Aspergillosis in 2019. Clin Microbiol Rev 33:e00140–18.

4. Liu K-W, Grau MS, Jones JT, Wang X, Vesely EM, James MR, Gutierrez-Perez C, Cramer RA, Obar JJ. 2022. Postinfluenza Environment Reduces Aspergillus fumigatus Conidium Clearance and Facilitates Invasive Aspergillosis In Vivo. mBio 13:e02854–22.

5. Sarden N, Sinha S, Potts KG, Pernet E, Hiroki CH, Hassanabad MF, Nguyen AP, Lou Y, Farias R, Winston BW, Bromley A, Snarr BD, Zucoloto AZ, Andonegui G, Muruve DA, McDonald B, Sheppard DC, Mahoney DJ, Divangahi M, Rosin N, Biernaskie J, Yipp BG. 2022. A B1a–natural IgG–neutrophil axis is impaired in viral-and steroid-associated aspergillosis. Science Translational Medicine 14:eabq6682.

6. Tio SY, Chen SC-A, Hamilton K, Heath CH, Pradhan A, Morris AJ, Korman TM, Morrissey O, Halliday CL, Kidd S, Spelman T, Brell N, McMullan B, Clark JE, Mitsakos K, Hardiman RP, Williams P, Campbell AJ, Beardsley J, Van Hal S, Yong MK, Worth LJ, Slavin MA. 2023. Invasive aspergillosis in adult patients in Australia and New Zealand: 2017–2020. The Lancet Regional Health - Western Pacific 40:100888.

7. WHO fungal priority pathogens list to guide research, development and public health action. https://www.who.int/publications/i/item/9789240060241. Retrieved 19 September 2025.

8. Patterson TF, Thompson GR, Denning DW, Fishman JA, Hadley S, Herbrecht R, Kontoyiannis DP, Marr KA, Morrison VA, Nguyen MH, Segal BH, Steinbach WJ, Stevens DA, Walsh TJ, Wingard JR, Young J-AH, Bennett JE. 2016. Practice Guidelines for the Diagnosis and Management of Aspergillosis: 2016 Update by the Infectious Diseases Society of America. Clin Infect Dis 63:e1–e60.

9. Verweij PE, Lucas JA, Arendrup MC, Bowyer P, Brinkmann AJF, Denning DW, Dyer PS, Fisher MC, Geenen PL, Gisi U, Hermann D, Hoogendijk A, Kiers E, Lagrou K, Melchers WJG, Rhodes J, Rietveld AG, Schoustra SE, Stenzel K, Zwaan BJ, Fraaije BA. 2020. The one health problem of azole resistance in *Aspergillus fumigatus*: current insights and future research agenda. Fungal Biology Reviews 34:202–214.

10. Lee Y, Robbins N, Cowen LE. 2023. Molecular mechanisms governing antifungal drug resistance. npj Antimicrob Resist 1:5.

11. Warrilow AGS, Parker JE, Price CL, Nes WD, Kelly SL, Kelly DE. 2015. In Vitro Biochemical Study of CYP51-Mediated Azole Resistance in Aspergillus fumigatus. Antimicrobial Agents and Chemotherapy 59:7771–7778.

12. Verweij PE, Lucas JA, Arendrup MC, Bowyer P, Brinkmann AJF, Denning DW, Dyer PS, Fisher MC, Geenen PL, Gisi U, Hermann D, Hoogendijk A, Kiers E, Lagrou K, Melchers WJG, Rhodes J, Rietveld AG, Schoustra SE, Stenzel K, Zwaan BJ, Fraaije BA. 2020. The one health problem of azole resistance in Aspergillus fumigatus: current insights and future research agenda. Fungal Biology Reviews 34:202–214.

13. Mellado E, Garcia-Effron G, Alcázar-Fuoli L, Melchers WJG, Verweij PE, Cuenca-Estrella M, Rodríguez-Tudela JL. 2007. A new Aspergillus fumigatus resistance mechanism conferring in vitro cross-resistance to azole antifungals involves a combination of cyp51A alterations. Antimicrob Agents Chemother 51:1897–1904.

14. Chowdhary A, Sharma C, Meis JF. 2017. Azole-Resistant Aspergillosis: Epidemiology, Molecular Mechanisms, and Treatment. J Infect Dis 216:S436–S444.

15. Gsaller F, Hortschansky P, Furukawa T, Carr PD, Rash B, Capilla J, Müller C, Bracher F, Bowyer P, Haas H, Brakhage AA, Bromley MJ. 2016. Sterol Biosynthesis and Azole Tolerance Is Governed by the Opposing Actions of SrbA and the CCAAT Binding Complex. PLOS Pathogens 12:e1005775.

16. Blosser SJ, Cramer RA. 2012. SREBP-Dependent Triazole Susceptibility in Aspergillus fumigatus Is Mediated through Direct Transcriptional Regulation of erg11A (cyp51A). Antimicrob Agents Chemother 56:248–257.

17. Blatzer M, Barker BM, Willger SD, Beckmann N, Blosser SJ, Cornish EJ, Mazurie A, Grahl N, Haas H, Cramer RA. 2011. SREBP coordinates iron and ergosterol homeostasis to mediate triazole drug and hypoxia responses in the human fungal pathogen Aspergillus fumigatus. PLoS Genet 7:e1002374.

18. Paul S, Stamnes M, Thomas GH, Liu H, Hagiwara D, Gomi K, Filler SG, Moye-Rowley WS. 2019. AtrR Is an Essential Determinant of Azole Resistance in Aspergillus fumigatus. mBio 10:10.1128/mbio.02563-18.

19. Willger SD, Puttikamonkul S, Kim K-H, Burritt JB, Grahl N, Metzler LJ, Barbuch R, Bard M, Lawrence CB, Jr RAC. 2008. A Sterol-Regulatory Element Binding Protein Is Required for Cell Polarity, Hypoxia Adaptation, Azole Drug Resistance, and Virulence in Aspergillus fumigatus. PLOS Pathogens 4:e1000200.

20. Yan R, Cao P, Song W, Li Y, Wang T, Qian H, Yan C, Yan N. 2021. Structural basis for sterol sensing by Scap and Insig. Cell Rep 35:109299.

21. Butler G. 2013. Hypoxia and Gene Expression in Eukaryotic Microbes. Annu Rev Microbiol 67:291–312.

22. Sever N, Yang T, Brown MS, Goldstein JL, DeBose-Boyd RA. 2003. Accelerated degradation of HMG CoA reductase mediated by binding of insig-1 to its sterol-sensing domain. Mol Cell 11:25–33.

23. Rybak JM, Ge W, Wiederhold NP, Parker JE, Kelly SL, Rogers PD, Fortwendel JR. 2019. Mutations in hmg1, Challenging the Paradigm of Clinical Triazole Resistance in Aspergillus fumigatus. mBio 10:10.1128/mbio.00437-19.

24. Hagiwara D, Arai T, Takahashi H, Kusuya Y, Watanabe A, Kamei K. 2018. Non-cyp51A Azole-Resistant Aspergillus fumigatus Isolates with Mutation in HMG-CoA Reductase. Emerg Infect Dis 24:1889–1897.

25. Gonzalez-Jimenez I, Lucio J, Roldan A, Alcazar-Fuoli L, Mellado E. 2021. Are Point Mutations in HMG-CoA Reductases (Hmg1 and Hmg2) a Step towards Azole Resistance in Aspergillus fumigatus? Molecules 26:5975.

26. Souza ACO, Ge W, Wiederhold NP, Rybak JM, Fortwendel JR, Rogers PD. 2023. hapE and hmg1 Mutations Are Drivers of cyp51A-Independent Pan-Triazole Resistance in an Aspergillus fumigatus Clinical Isolate. Microbiology Spectrum 11:e05188–22.

27. Rybak JM, Xie J, Martin-Vicente A, Guruceaga X, Thorn HI, Nywening AV, Ge W, Souza ACO, Shetty AC, McCracken C, Bruno VM, Parker JE, Kelly SL, Snell HM, Cuomo CA, Rogers PD, Fortwendel JR. 2024. A secondary mechanism of action for triazole antifungals in Aspergillus fumigatus mediated by hmg1. Nat Commun 15:3642.

28. Berman J, Krysan DJ. 2020. Drug resistance and tolerance in fungi. Nat Rev Microbiol 18:319–331.

29. Scott J, Valero C, Mato-López Á, Donaldson IJ, Roldán A, Chown H, Rhijn NV, Lobo-Vega R, Gago S, Furukawa T, Morogovsky A, Ami RB, Bowyer P, Osherov N, Fontaine T, Goldman GH, Mellado E, Bromley M, Amich J. 2022. Aspergillus fumigatus can display persistence to the fungicidal drug voriconazole. bioRxiv 10.1101/2022.05.16.491816.

30. Scott J, Valero C, Mato-López Á, Donaldson IJ, Roldán A, Chown H, Rhijn NV, Lobo-Vega R, Gago S, Furukawa T, Morogovsky A, Ami RB, Bowyer P, Osherov N, Fontaine T, Goldman GH, Mellado E, Bromley M, Amich J. 2023. Aspergillus fumigatus Can Display Persistence to the Fungicidal Drug Voriconazole. Microbiology Spectrum 10.1128/spectrum.04770-22.

31. Amich J, Bromley M, Goldman GH, Valero C. 2025. Toward the consensus of definitions for the phenomena of antifungal tolerance and persistence in filamentous fungi. mBio 16:e03475–24.

32. Basenko EY, Shanmugasundram A, Böhme U, Starns D, Wilkinson PA, Davison HR, Crouch K, Maslen G, Harb OS, Amos B, McDowell MA, Kissinger JC, Roos DS, Jones A. 2024. What is new in FungiDB: a web-based bioinformatics platform for omics-scale data analysis for fungal and oomycete species. Genetics 227:iyae035.

33. Jöchl C, Rederstorff M, Hertel J, Stadler PF, Hofacker IL, Schrettl M, Haas H, Hüttenhofer A. 2008. Small ncRNA transcriptome analysis from Aspergillus fumigatus suggests a novel mechanism for regulation of protein synthesis. Nucleic Acids Res 36:2677–2689.

34. Scotto–Lavino E, Du G, Frohman MA. 2006. 3′ End cDNA amplification using classic RACE. Nat Protoc 1:2742–2745.

35. Kang Y-J, Yang D-C, Kong L, Hou M, Meng Y-Q, Wei L, Gao G. 2017. CPC2: a fast and accurate coding potential calculator based on sequence intrinsic features. Nucleic Acids Res 45:W12–W16.

36. Snarr BD, Baker P, Bamford NC, Sato Y, Liu H, Lehoux M, Gravelat FN, Ostapska H, Baistrocchi SR, Cerone RP, Filler EE, Parsek MR, Filler SG, Howell PL, Sheppard DC. 2017. Microbial glycoside hydrolases as antibiofilm agents with cross-kingdom activity. Proc Natl Acad Sci U S A 114:7124–7129.

37. Bertuzzi M, van Rhijn N, Krappmann S, Bowyer P, Bromley MJ, Bignell EM. 2020. On the lineage of Aspergillus fumigatus isolates in common laboratory use. Med Mycol 59:7–13.

38. Keller NP. 2017. Heterogeneity Confounds Establishment of “a” Model Microbial Strain. mBio 8:10.1128/mbio.00135-17.

39. Dhingra S, Kowalski CH, Thammahong A, Beattie SR, Bultman KM, Cramer RA. 2016. RbdB, a Rhomboid Protease Critical for SREBP Activation and Virulence in Aspergillus fumigatus. mSphere 1:e00035–16.

40. Dhingra S, Buckey JC, Cramer RA. 2018. Hyperbaric Oxygen Reduces Aspergillus fumigatus Proliferation In Vitro and Influences In Vivo Disease Outcomes. Antimicrob Agents Chemother 62:e01953–17.

41. Graybill JR, Najvar LK, Gonzalez GM, Hernandez S, Bocanegra R. 2003. Improving the mouse model for studying the efficacy of voriconazole. J Antimicrob Chemother 51:1373–1376.

42. Manente M, Ghislain M. 2009. The lipid-translocating exporter family and membrane phospholipid homeostasis in yeast. FEMS Yeast Res 9:673–687.

43. Hokken MWJ, Zoll J, Coolen JPM, Zwaan BJ, Verweij PE, Melchers WJG. 2019. Phenotypic plasticity and the evolution of azole resistance in Aspergillus fumigatus; an expression profile of clinical isolates upon exposure to itraconazole. BMC Genomics 20:28.

44. Ness F, Aigle M. 1995. Rtm1: A Member of a New Family of Telomeric Repeated Genes in Yeast. Genetics 140:945–956.

45. Abou-Kandil A, Tröger-Görler S, Pschibul A, Krüger T, Rosin M, Schmidt F, Akbarimoghaddam P, Sarkar A, Cseresnyés Z, Shadkchan Y, Heinekamp T, Gräler MH, Barber AE, Walther G, Figge MT, Brakhage AA, Osherov N, Kniemeyer O. 2025. The proteomic response of Aspergillus fumigatus to amphotericin B (AmB) reveals the involvement of the RTA-like protein RtaA in AmB resistance. microLife 6:uqae024.

46. Verweij PE, Chowdhary A, Melchers WJG, Meis JF. 2016. Azole Resistance in Aspergillus fumigatus: Can We Retain the Clinical Use of Mold-Active Antifungal Azoles? Clin Infect Dis 62:362–368.

47. Cuypers L, Aerts R, Van de gaer O, Vinken L, Merckx R, Gerils V, Vande Velde G, Reséndiz-Sharpe A, Maertens J, Lagrou K. 2025. Doubling of triazole resistance rates in invasive aspergillosis over a 10-year period, Belgium, 1 April 2022 to 31 March 2023. Euro Surveill 30:2400559.

48. Kang Y, Ma W, Li Q, Wang P, Jia W. 2025. Epidemiology, antifungal susceptibility and biological characteristics of clinical Aspergillus fumigatus in a tertiary hospital. Sci Rep 15:16906.

49. Burks C, Darby A, Londoño LG, Momany M, Brewer MT. 2021. Azole-resistant Aspergillus fumigatus in the environment: Identifying key reservoirs and hotspots of antifungal resistance. PLOS Pathogens 17:e1009711.

50. Liu S, Le Mauff F, Sheppard DC, Zhang S. 2022. Filamentous fungal biofilms: Conserved and unique aspects of extracellular matrix composition, mechanisms of drug resistance and regulatory networks in Aspergillus fumigatus. npj Biofilms Microbiomes 8:83.

51. Kowalski CH, Morelli KA, Schultz D, Nadell CD, Cramer RA. 2020. Fungal biofilm architecture produces hypoxic microenvironments that drive antifungal resistance. Proceedings of the National Academy of Sciences 117:22473–22483.

52. Kowalski CH, Morelli KA, Stajich JE, Nadell CD, Cramer RA. 2021. A Heterogeneously Expressed Gene Family Modulates the Biofilm Architecture and Hypoxic Growth of Aspergillus fumigatus. mBio 12:e03579–20.

53. Wuyts J, Van Dijck P, Holtappels M. 2018. Fungal persister cells: The basis for recalcitrant infections? PLoS Pathog 14:e1007301.

54. De Francesco MA. 2023. Drug-Resistant Aspergillus spp.: A Literature Review of Its Resistance Mechanisms and Its Prevalence in Europe. Pathogens 12:1305.

55. Dagenais TRT, Keller NP. 2009. Pathogenesis of Aspergillus fumigatus in Invasive Aspergillosis. Clin Microbiol Rev 22:447–465.

56. Bueid A, Howard SJ, Moore CB, Richardson MD, Harrison E, Bowyer P, Denning DW. 2010. Azole antifungal resistance in Aspergillus fumigatus: 2008 and 2009. J Antimicrob Chemother 65:2116–2118.

57. Zhao Y, Prideaux B, Baistrocchi S, Sheppard DC, Perlin DS. 2019. Beyond tissue concentrations: antifungal penetration at the site of infection. Med Mycol 57:S161–S167.

58. Moreillon P, Tomasz A. 1988. Penicillin resistance and defective lysis in clinical isolates of pneumococci: evidence for two kinds of antibiotic pressure operating in the clinical environment. J Infect Dis 157:1150–1157.

59. Windels EM, Michiels JE, Van den Bergh B, Fauvart M, Michiels J. 2019. Antibiotics: Combatting Tolerance To Stop Resistance. mBio 10:e02095–19.

60. Conte JE, Golden JA, Krishna G, McIver M, Little E, Zurlinden E. 2009. Intrapulmonary Pharmacokinetics and Pharmacodynamics of Posaconazole at Steady State in Healthy Subjects. Antimicrob Agents Chemother 53:703–707.

61. Campoli P, Al Abdallah Q, Robitaille R, Solis NV, Fielhaber JA, Kristof AS, Laverdiere M, Filler SG, Sheppard DC. 2011. Concentration of Antifungal Agents within Host Cell Membranes: a New Paradigm Governing the Efficacy of Prophylaxis. Antimicrob Agents Chemother 55:5732–5739.

62. Kowalski CH, Beattie SR, Fuller KK, McGurk EA, Tang Y-W, Hohl TM, Obar JJ, Cramer RA. 2016. Heterogeneity among Isolates Reveals that Fitness in Low Oxygen Correlates with Aspergillus fumigatus Virulence. mBio 7:10.1128/mbio.01515-16.

63. Caffrey-Carr AK, Kowalski CH, Beattie SR, Blaseg NA, Upshaw CR, Thammahong A, Lust HE, Tang Y-W, Hohl TM, Cramer RA, Obar JJ. 2017. Interleukin 1α Is Critical for Resistance against Highly Virulent Aspergillus fumigatus Isolates. Infect Immun 85:e00661–17.

64. Fuller KK, Cramer RA, Zegans ME, Dunlap JC, Loros JJ. 2016. Aspergillus fumigatus Photobiology Illuminates the Marked Heterogeneity between Isolates. mBio 7:e01517–16.

65. Traubenik S, Charon C, Blein T. 2024. From environmental responses to adaptation: the roles of plant lncRNAs. Plant Physiol 195:232–244.

66. Bowyer P, Weaver D, Qi T, Chown H, Fraczek M, Lebedinec R, Dineen L, Valero C, Rhijn N van, Furukawa T, Bromley M, Delneri D. 2025. Genome-wide discovery and phenotyping of non-coding transcripts in A. fumigatus reveals lncRNAs with a role in antifungal drug sensitivity. Research Square 10.21203/rs.3.rs-6048166/v1.

67. Smith-Peavler ES, Patel R, Onumajuru AM, Bowring BG, Miller JL, Brunel JM, Djordjevic JT, Prabu MM, McClelland EE. 2022. RTA1 Is Involved in Resistance to 7-Aminocholesterol and Secretion of Fungal Proteins in Cryptococcus neoformans. Pathogens 11:1239.

68. Barber AE, Sae-Ong T, Kang K, Seelbinder B, Li J, Walther G, Panagiotou G, Kurzai O. 2021. Aspergillus fumigatus pan-genome analysis identifies genetic variants associated with human infection. Nat Microbiol 6:1526–1536.

69. Mercer TR, Dinger ME, Mattick JS. 2009. Long non-coding RNAs: insights into functions. Nat Rev Genet 10:155–159.

70. Thammahong A, Caffrey-Card AK, Dhingra S, Obar JJ, Cramer RA. 2017. Aspergillus fumigatus Trehalose-Regulatory Subunit Homolog Moonlights To Mediate Cell Wall Homeostasis through Modulation of Chitin Synthase Activity. mBio 8:e00056–17.

71. Szewczyk E, Nayak T, Oakley CE, Edgerton H, Xiong Y, Taheri-Talesh N, Osmani SA, Oakley BR. 2006. Fusion PCR and gene targeting in Aspergillus nidulans. Nat Protoc 1:3111–3120.

72. Thammahong A, Dhingra S, Bultman KM, Kerkaert JD, Cramer RA. 2019. An Ssd1 Homolog Impacts Trehalose and Chitin Biosynthesis and Contributes to Virulence in Aspergillus fumigatus. mSphere 4:e00244–19.

73. Al Abdallah Q, Ge W, Fortwendel JR. 2017. A Simple and Universal System for Gene Manipulation in Aspergillus fumigatus: In Vitro-Assembled Cas9-Guide RNA Ribonucleoproteins Coupled with Microhomology Repair Templates. mSphere 2:e00446–17.

74. Furukawa T, van Rhijn N, Chown H, Rhodes J, Alfuraji N, Fortune-Grant R, Bignell E, Fisher MC, Bromley M. 2022. Exploring a novel genomic safe-haven site in the human pathogenic mould Aspergillus fumigatus. Fungal Genet Biol 161:103702.

75. Oakley BR, Rinehart JE, Mitchell BL, Oakley CE, Carmona C, Gray GL, May GS. 1987. Cloning, mapping and molecular analysis of the pyrG (orotidine-5’-phosphate decarboxylase) gene of Aspergillus nidulans. Gene 61:385–399.

76. Dhingra S, Lind AL, Lin H-C, Tang Y, Rokas A, Calvo AM. 2013. The fumagillin gene cluster, an example of hundreds of genes under veA control in Aspergillus fumigatus. PLoS One 8:e77147.

77. Osmani AH, May GS, Osmani SA. 1999. The extremely conserved pyroA gene of Aspergillus nidulans is required for pyridoxine synthesis and is required indirectly for resistance to photosensitizers. J Biol Chem 274:23565–23569.

78. Zepeda B, Verdonk JC. 2022. RNA Extraction from Plant Tissue with Homemade Acid Guanidinium Thiocyanate Phenol Chloroform (AGPC). Current Protocols 2:e351.

79. STAR: ultrafast universal RNA-seq aligner | Bioinformatics | Oxford Academic. https://academic.oup.com/bioinformatics/article/29/1/15/272537. Retrieved 9 September 2025.

80. Danecek P, Bonfield JK, Liddle J, Marshall J, Ohan V, Pollard MO, Whitwham A, Keane T, McCarthy SA, Davies RM, Li H. 2021. Twelve years of SAMtools and BCFtools. Gigascience 10:giab008.

81. Liao Y, Smyth GK, Shi W. 2014. featureCounts: an efficient general purpose program for assigning sequence reads to genomic features. Bioinformatics 30:923–930.

82. Love MI, Huber W, Anders S. 2014. Moderated estimation of fold change and dispersion for RNA-seq data with DESeq2. Genome Biol 15:550.

83. Blighe K, Rana S, Lewis M. EnhancedVolcano: Publication-ready volcano plots with enhanced colouring and labeling 10.18129/B9.bioc.EnhancedVolcano.

84. R Core Team (2024). R: A Language and Environment for Statistical Computing (version 3.5.1). R Foundation for Statistical Computing. https://www.R-project.org/. Retrieved 9 September 2025.

85. Stajich J. 2025. stajichlab/DhingraLab_ncRNA_Afumigatus: Archive of Analysis and Data archive for publication (v0.1). Zenodo.

