## Supplementary file for "A long non-coding RNA regulates in vitro and in vivo triazole antifungal susceptibility in *Aspergillus fumigatus*"

Figure S1:

A

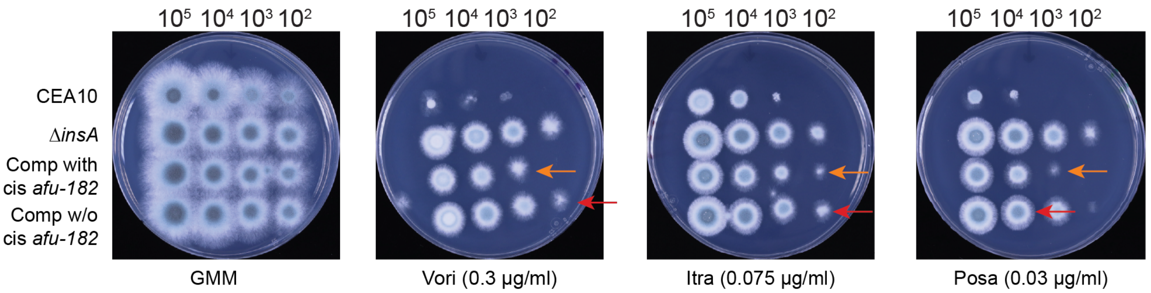


B C

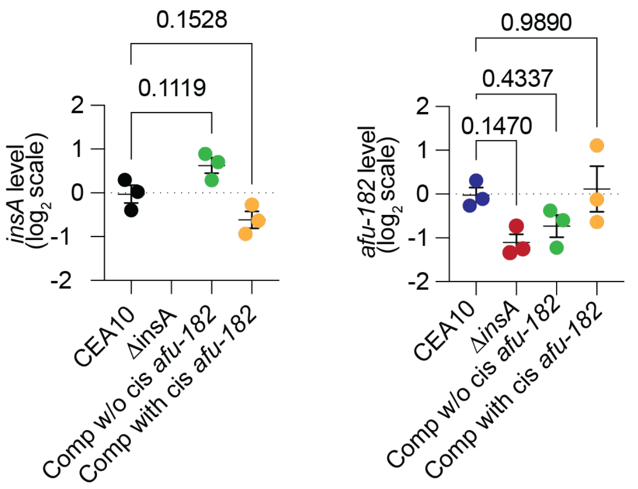


D

| Strain | Voriconazole MIC (μg/ml) |
|---|---|
| CEA10 | 0.25-0.5 |
| Δ*insA* | 0.25-0.5 |
| Comp with cis *afu-182* | 0.25-0.5 |
| Comp w/o cis *afu-182* | 0.25-0.5 |

**Figure S1. *insA* regulates fungal azole response.** (A) AF strains were serially inoculated ($10^5$ – $10^2$) on GMM in the absence and presence of azole drugs as indicated. The Δ*insA* strain showed increased fungal growth in the presence of the sub-MIC concentration of azoles; however, complementation of *insA* without cis-acting *afu-182* does not revert the phenotype back to WT (red arrows), whereas complementation of *insA* with cis-acting *afu-182* partially reverts the phenotype back to WT levels (orange arrows). (B and C) Quantification of *insA* and *afu-182* RNA levels. Quantitative reverse-transcription PCR was used to assess the RNA levels of (B) *insA* and (C) *afu-182*. (D) The minimum inhibitory concentration of the indicated strains was measured via broth microdilution assay. No change in MIC was observed for voriconazole.

Figure S2:

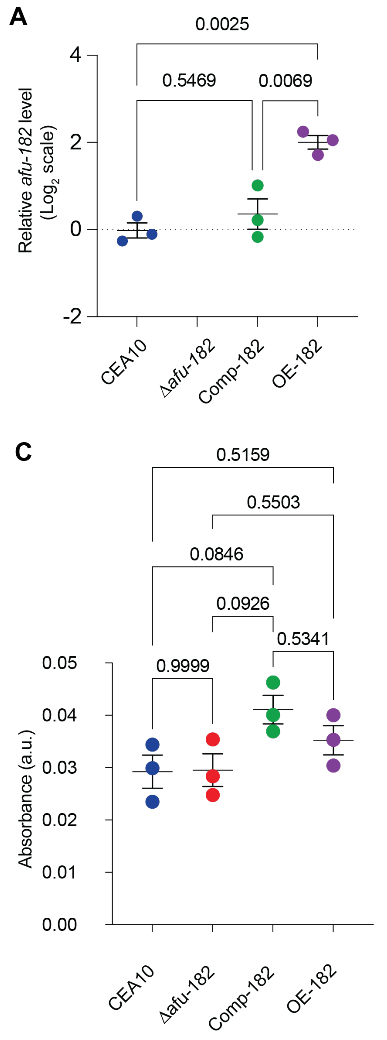


B

| Strain | Voriconazole MIC (μg/ml) |
|---|---|
| CEA10 | 0.25-0.5 |
| *Δafu-182* | 0.25-0.5 |
| Comp-182 | 0.25-0.5 |
| OE-182 | 0.25-0.5 |

**Figure S2.** *afu-182* affects sub-MIC azole growth without a change in MIC. (A) Quantitative reverse-transcription PCR was used to assess the RNA levels of *afu-182* in WT, *Δafu-182*, Comp-182 and OE-182 strains. No transcript was detected in *Δafu-182* and 4-fold overexpression was observed in the OE-182 strain (p=0.0025, One-Way ANOVA). (B) The table highlights the minimum inhibitory concentration of voriconazole using the broth microdilution method. No change in MIC was observed. (C) Effects of *afu-182* on fungal surface attachment. AF strains were allowed to adhere to cell culture plates for 12 hours before azole treatment and stained with crystal violet. The $OD_{600}$ values are represented. No difference in surface attachment was observed. a.u. – arbitrary units.

Figure S3:

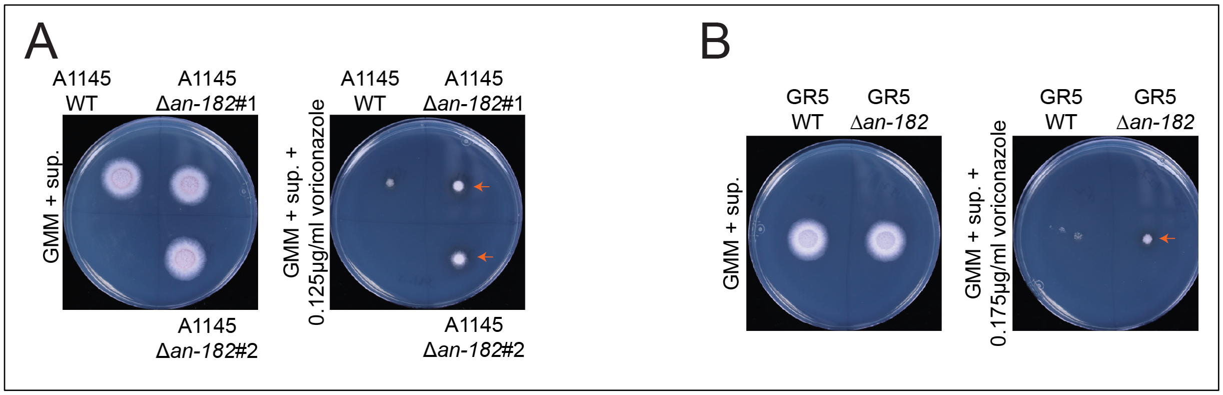


**Figure S3. *afu-182* regulates fungal sub-MIC growth in *A. nidulans*.** *afu-182* homolog (*an-182*) was identified in the related fungal species *Aspergillus nidulans*. *an-182* was deleted in A1145 or GR5 strains of AN. $10^4$ spores of the indicated strains were spot inoculated on GMM with supplements (see methods) in the absence and presence of voriconazole. Increased fungal growth at sub-MIC azole concentration was observed in (A) A1145 and (B) GR5 strains (orange arrows). #1 and #2 indicate two independent deletion transformants.

Figure S4:

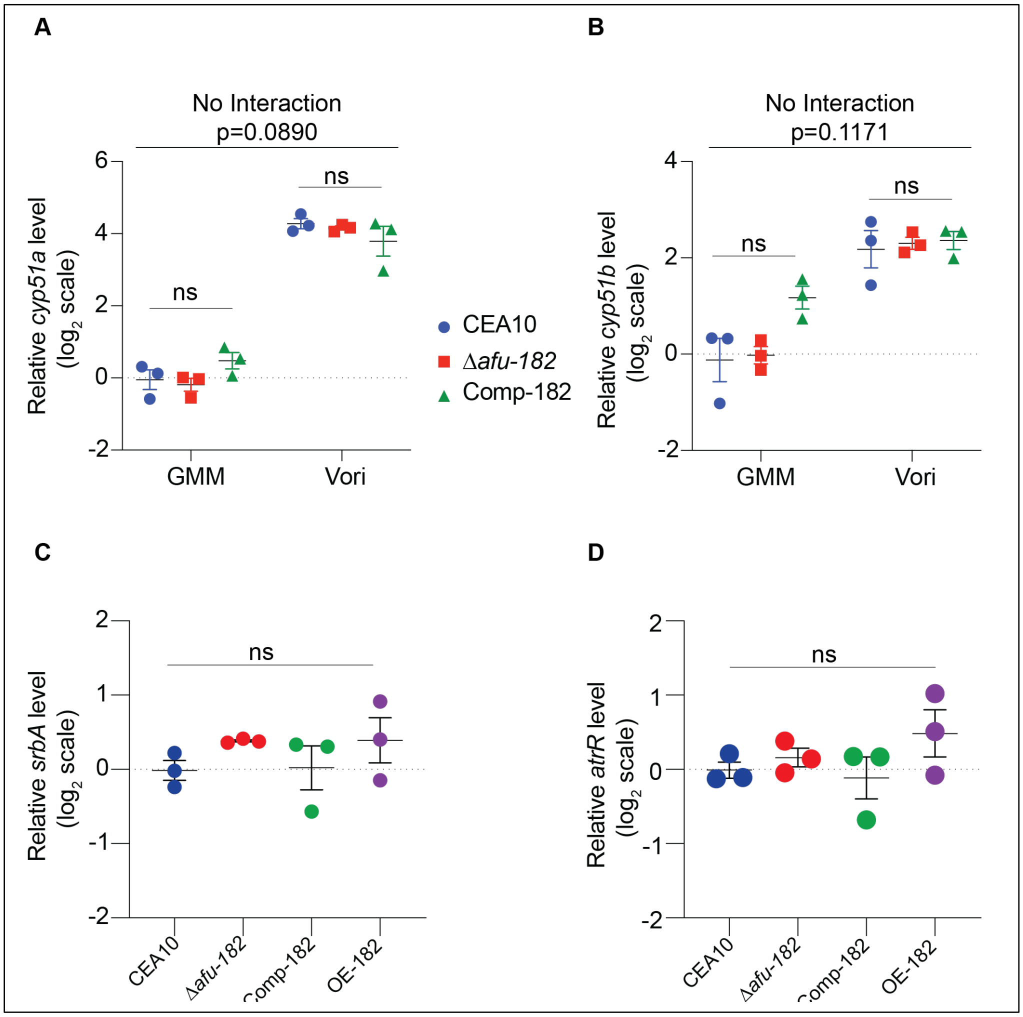


**Figure S4. *afu-182* regulates fungal azole response independent of *cyp51* transcriptional regulation**. Quantitative reverse transcriptase was used to determine the mRNA levels of (A) *cyp51a* and (B) *cyp51b* in the presence or absence of azole drugs. No significant difference due to genotype was observed for *cyp51a* (interaction p=0.3291, Two-Way ANOVA) or *cyp51b* (interaction p=0.1171, Two-Way ANOVA). mRNA levels of (C) *srbA* and (D) *atrR* were similarly quantified in GMM. No significant difference was observed between WT and Δ*afu-182* strains (One-Way ANOVA followed by Tukey's post hoc analysis). ns – not significant

Figure S5:

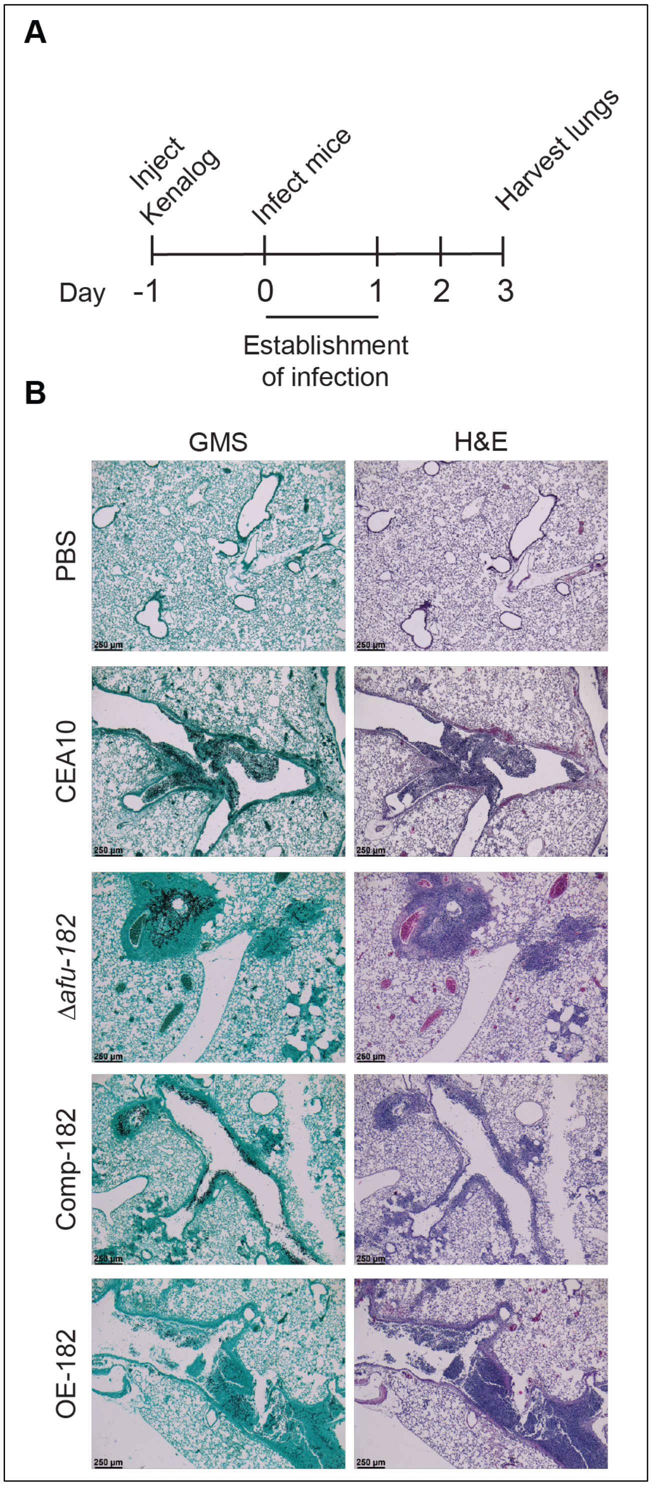


**Figure S5.** (A) Schematic showing animal experimental protocol. (B) Histopathological analysis of murine lungs harvested as in A. No qualitative differences were seen.

Figure S6:

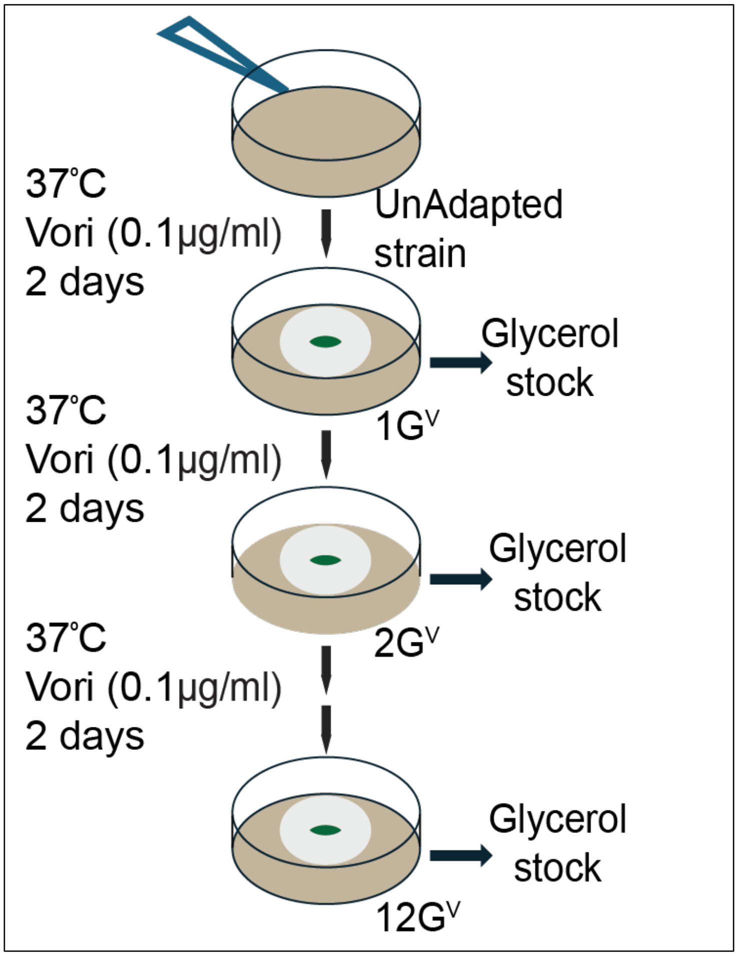


**Figure S6.** A schematic showing azole adaptation experiment for WT and Δ*afu-182* strain.

Figure S7:

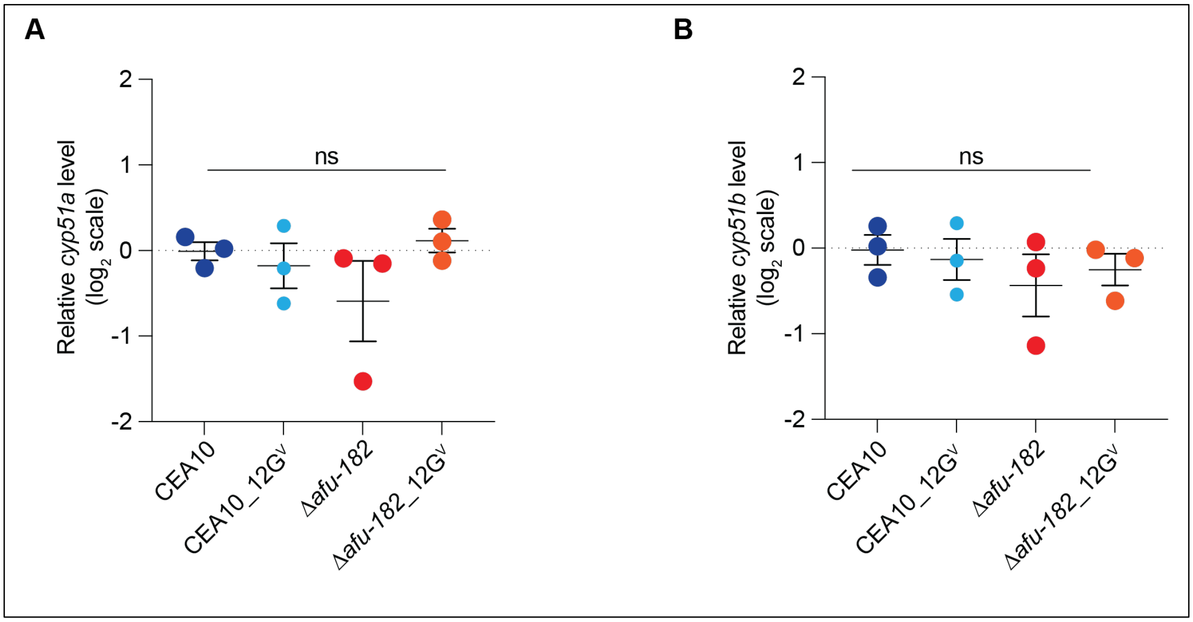


**Figure S7. *afu-182* mediated increase in azole MIC is independent of *cyp51* transcriptional regulation**. Quantitative reverse transcriptase was used to determine the mRNA levels of (A) *cyp51a* and (B) *cyp51b*. No significant difference was observed (One-Way ANOVA followed by Tukey's post hoc analysis). ns – not significant

Figure S8:

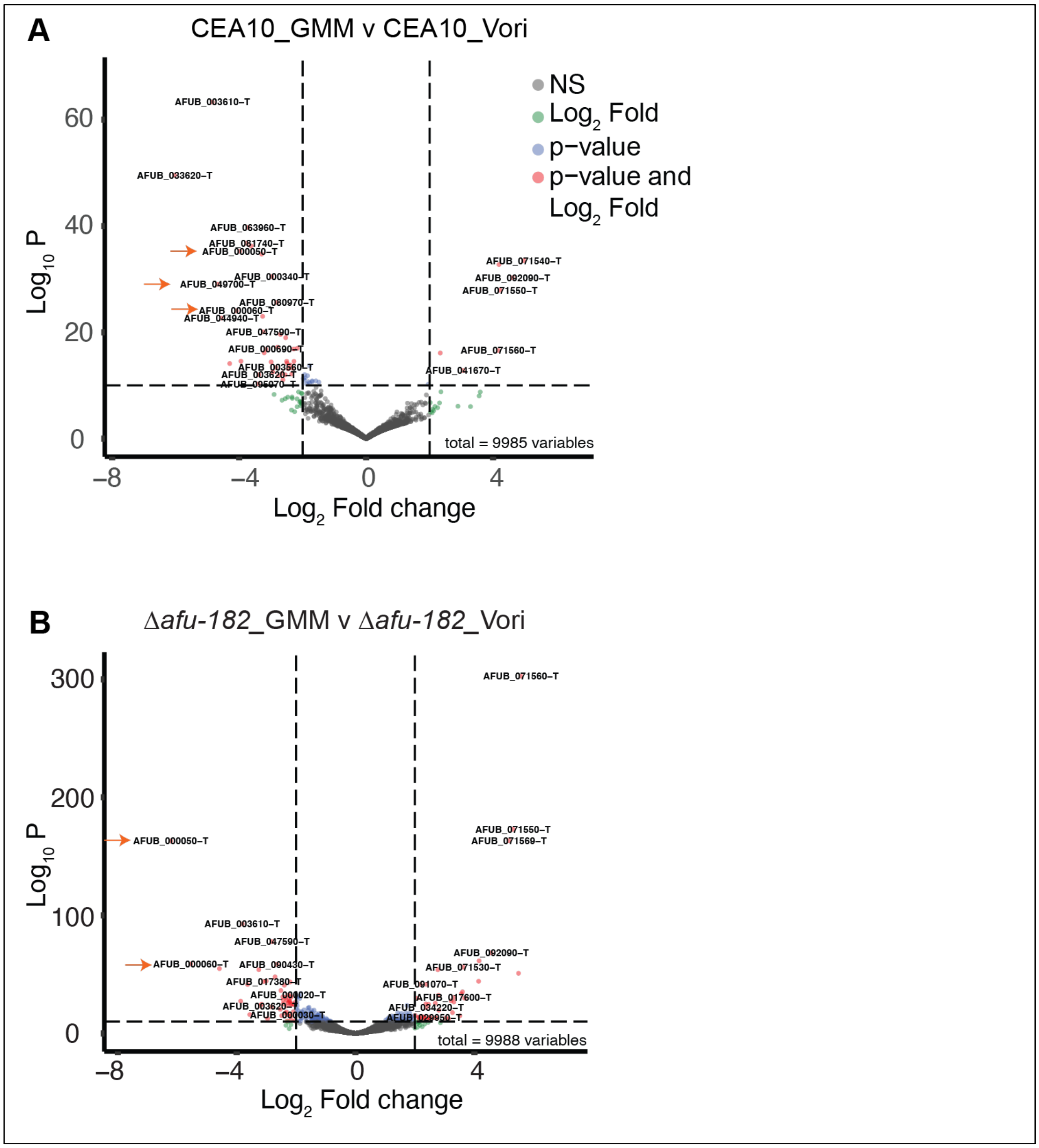


**Figure S8.** RTA1 domain proteins are upregulated in presence of azole drugs. Volcano plots showing the 2-$log_2$ fold change (4-fold upregulated or downregulated). (A) Comparison between CEA10 GMM and CEA10 azole. Global analysis revealed genes encoding RTM1 domain proteins *rtmA* (*afub_000050,*) and *rtmB* (*afub_000060*) and afub_049700 (orange arrows) are significantly upregulated. (B) Comparison between Δ*afu-182* GMM and Δ*afu-182* azoles. *rtmA* and *rtmB* genes are significantly upregulated in the presence of azoles in the Δ*afu-182* strain compared to the GMM control.

Figure S9:

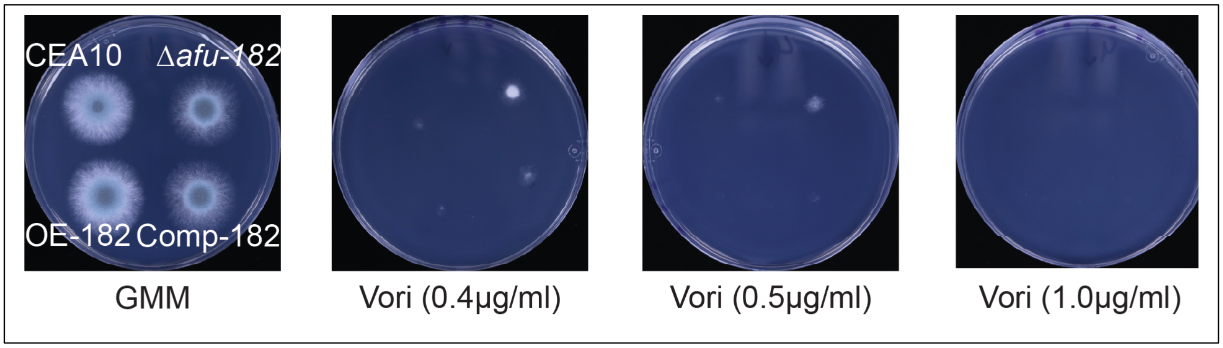


**Figure S3. *afu-182* does not regulate growth at supra-MIC.** AF strains were inoculated on GMM in the absence and presence of voriconazole approaching MIC (0.4μg/ml), MIC (0.5μg/ml) and 2x MIC (1.0 μg/ml). Robust growth was observed in Δafu-182 strain at 0.4μg/ml. Germination was observed for all strains at 0.5μg/ml and no growth was observed at 1μg/ml for all strains after 48 hours.

Figure S10:

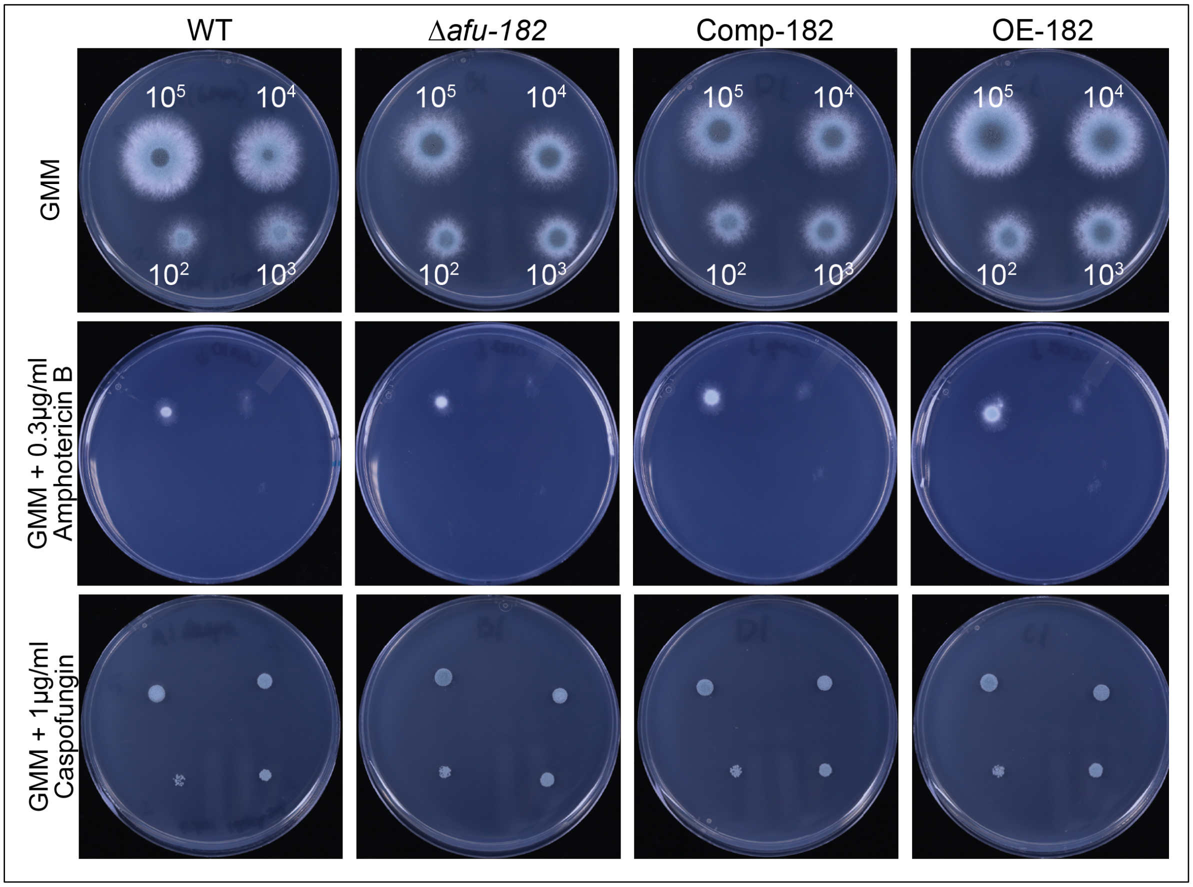


**Figure S10. *afu-182* is dispensable for fungal response to amphotericin B and Caspofungin.** AF strains were serially inoculated ($10^5 - 10^2$) on GMM in the absence and presence of 0.3μg/ml amphotericin B or 1μg/ml Caspofungin. No difference was observed.

**Table S1. List of strains used in this study:**

| Strain | Genotype | Source |
|---|---|---|
| CEA10 | WT | FGSC |
| CEA17 | *pyrG-* | FGSC |
| Δ*insA* | *pyrG-*; Δ*insA*::*pyrG A. parasiticus* | This study |
| Comp-insA with cis-acting *afu-182* | *pyrG-*; Δ*insA*::*pyrG A. parasiticus*; *insA*(P)::*insA*::*afu-182*::*insA*(T); Hygromycin+ | This study |
| Comp-insA without cis-acting *afu-182* | *pyrG-*; Δ*insA*::*pyrG A. parasiticus*; *insA*(P)::*insA*::*insA*(T); Hygromycin+ | This study |
| Δ*afu-182* | *pyrG-*; Δ*afu-182*::*pyrG A. parasiticus* | This study |
| Comp-182 | *pyrG-*; Δ*afu-182*::*pyrG A. parasiticus*; *afu-182*::*ptrA* | This study |
| OE *afu-182* | *pyrG-*; *gpdA*(+1TSS)::*afu-182*::*trpC*(T), *pyrG* | This study |
| A1145 | *pyrG89*; *pyroA4*;*nkuA*::*argB*; *riboB2* | FGSC |
| A1145 Δan-182 | *pyrG89*; *pyroA4*;*nkuA*::*argB*; *riboB2*; Δ*an-182*::*pyrG* | This study |
| GR5 | *pyrG89*; *wA3*; *pyroA4*; | FGSC |
| GR5 Δan-182 | *pyrG89*; *wA3*; *pyroA4*; Δan-182::*pyroA A. fumigatus* | This study |
| ΔrtmA | Δ*rtmA*::*ptrA* | This study |
| ΔrtmB | Δ*rtmB*::*ptrA* | This study |
| ΔrtmAΔrtmB | Δ*rtmA*Δ*rtmB*::*ptrA* | This study |
| Comp-rtmA | Δ*rtmA*::*ptrA*; *rtmA*(p)::*rtmA*::*rtmA*(T):; Hygromycin | This study |

| | | |
|---|---|---|
| Comp-rtmB | Δ*rtmB*::*ptrA*; *rtmB*(p)::*rtmB*::*rtmB*(T)::; Hygromycin | This study |
| Af293 | WT | FGSC |
| Af293.1 | *pyrG-* | FGSC |
| Af293 Δ*afu-182* | *pyrG-*,Δ*afu-182*::*pyrG A. parasiticus* | This study |
| 47-57 | Prototroph | |
| 47-57 Δ*afu-182* | Δ*afu-182*:ptrA+ | This study |

**Table S2. List of Primers used in the study:**

| Primer Name | Sequence (5' -> 3') |
|---|---|
| RAC 815 | TGTGGTGCAACTCAAGTGGAGAGA |
| RAC 816 | CCAACTTAATCGCCTTGCAGCACAATCAGGTAGTTGCGGTCTCCCAAA |
| RAC819 | ATTCCACACAACATACGAGCCGGAAGGCCGAAGCTTTGGGTATGATCT |
| RAC820 | AAGCGTCACCAGTCTCACTGTCAA |
| RAC817 | TTTGGGAGACCGCAACTACCTGATTGTGCTGCAAGGCGATTAAGTTGG |
| RAC818 | AGATCATACCCAAAGCTTCGGCCTTCCGGCTCGTATGTTGTGTGGAAT |
| RAC 821 | GGGGACAAGTTTGTACAAAAAAGCAGGCTGAGCATCGTAGAAAGTCCCACGGGAAA |
| RAC 822 | GGGGACCACTTTGTACAAGAAAGCTGGGTTTTCTATCTCTGCTCATCCCACCG |
| SD37_linker | cggcgcgccagatctacgcgtttaattaaCCGC |
| SD38_linker | GGttaattaaacgcgtagatctggcgcgccGAGCT |
| RAC3589 | AAAAAAGGCGCGCCCAGCCTTGGAGCTGTCGAGTG |
| RAC3591 | AAAAAATTAATTAACACATGACACAGCACAGACCACC |
| RAC3590 | AAAAAATTAATTAAACATCGTTGACATGGCAAAGTCCG |

| | |
|---|---|
| SD149_182 homology F | GACCACTTCGACAACACACGACTAGACACCATACAACCGGTCGCCTCAAACAATGCTCT |
| SD150_182 homology | AAAACTTCTTTCCTTTTTCTTTTCTTCTTTTTCCTGTCTGAGAGGAGGCACTGATGCG |
| SD76 | CAGGCGTCTGCCACTCTTGC |
| SD164 | ccgtctgtcagatccccagagCGTGAGACACTTGACAGCGATGG |
| SD23 | cgactcactatagggagagcggc |
| SD165 | ctctggggatctgacagacgg |
| SD183 | GACCACTTCGACAACACACGACTAGACACCATACA |
| SD166 | tttctggtatattgttctgagatccataggatccaGTCTGAGAGGAGGCACTGATGCG |
| SD134 | CGAAGGCTTGGGGCACCTG |
| SD103 | GGGAAAAGAAAGAGAAAAGAAAAGAGCA |
| SD104 | TGCTCTTTTCTTTTCTCTTTCTTTTCCCTCATGAACAACGCAATTAACAACATTC |
| SD105 | GCGTTTTATTCTTGTTGACATGGGCTGTAGAGCCAATCATTGCAGT |
| SD135_EcoRI | NNNNNNGAATTCCcgagctcccaaatctgtccagatc |
| SD139_NotI | NNNNNNGCGGCCGCGGCTGTAGAGCCAATCATTGCAGT |
| SD162_OE_homology-F | TGCATCTTCCAGTTCTGGATATAGCATATAGATCTCCTAGCTGATTCTGGAGTGACCC |

| | |
|---|---|
| SD163_OE_homology-R | CCTCCATACTCCCCTGATCTCAATCCAGTTGAGAAGAGGAGGCACTGATGCGTGATG |
| SD181 | CGCTACAGGGTGATTCATAGAATCACAGCTTTCCAaccggtcgcctcaaacaatgctct |
| SD182 | AATAGCATCATCACGTGAAATAACGTGATGCACCGgtctgagaggaggcactgatgcg |
| SD 352 | CGCTACAGGGTGATTCATAGAATCACAGCTTTCCATGCCCATATCTTCCGTAGCAGTC |
| SD 353 | AATAGCATCATCACGTGAAATAACGTGATGCACCGTGACGGAGAGCTGAGAGTCCTAG |
| RAC2939 | ACCGGTCGCCTCAAACAATGCTCTATCCTTGAAGCTGTCCCTGATGGT |
| RAC2940 | GTCTGAGAGGAGGCACTGATGCGCGGCGTATTGGGTGTTACGGAGC |
| SD738 | NNNNNNNNGGCGCGCCCGTGAGCAACAGTGAACAATGCGAG |
| SD 739 | NNNNNNNNTTAATTAAGGAAGGTAAGGTGGCTGGGAC |
| SD740 | NNNNNNNNGGCGCGCCGGTGAGAGCAACTGTAAACGACTGAC |
| SD 741 | NNNNNNNNTTAATTAAGACCTGAACCTGTTCGTTGGTGTG |
| SD151 | TATTGCCCCTCTTGCATTCTTTATTGAAATTCTCCaccggtcgcctcaaacaatgctct |
| SD152 | CACAGAGTAGAGGAAGTAGACACGCCTTATTGCCTGTCTGAGAGGAGGCACTGATGCG |

| | |
|---|---|
| SD153 | CCTCTTCCTTCTCACGGCCATGTGTCCTGGCTCAAACCGGTCGCCTCAAACAATGCTCT |
| SD154 | GATAACTGTTCCTAGTCTTCGAAGCAAACATTCCTGTCTGAGAGGAGGCACTGATGCG |
| SD738 | NNNNNNNNGGCGCGCCCGTGAGCAACAGTGAACAATGCGAG |
| SD 739 | NNNNNNNNTTAATTAAGGAAGGTAAGGTGGCTGGGAC |
| SD740 | NNNNNNNNGGCGCGCCGGTGAGAGCAACTGTAAACGACTGAC |
| SD 741 | NNNNNNNNTTAATTAAGACCTGAACCTGTTCGTTGGTGTG |
| SD750_049700_qPCR F | GAACTGCAAGAGGGTTTCGACGG |
| SD751_049700_qPCR R | CTGGTCTGATCCCAGTAGCCACC |
| SD756_046760_qPCR F | GAGTGGTGCCTGTATGTCTTTGACG |
| SD757_046760_qPCR R | GGCAGCCGAAATCCCAGATATGG |
| SD758_000610_qPCR F | GGTCATTGTCGGCGAGGTCTTC |
| SD759_000610_qPCR R | GCCATGTTTAGTGTCCTGCAAGAGG |
| SD760_038360_qPCR F | GGACCATCGAGCTATCCGAAGG |
| SD761_038360_qPCR R | CGGTGGGCATCATCCAGCC |
| SD762_047370_qPCR F | GGGCTGTTCATGGTCACGTCG |
| SD763_047370_qPCR R | CTCGATCACCCTAAACACCGACC |
| SD764_088540_qPCR F | GGCGGGGATCTTGGTCACG |
| SD765_088540_qPCR R | GCAGCAATGAACCATCCCAGCC |
| SD766_053180_qPCR F | GGTTGATTTCTTCAGCCTCCTCCTCC |

| | |
|---|---|
| SD767_053180_qPCR R | CCCATTCATCAACTCCATGCTCCG |
| SD768_026620_qPCR F | CGTGCTCGGGGACATCTTGTC |
| SD769_026620_qPCR R | CATTATGGCACTGTGGTGTCGGG |
| SD772_036340_qPCR F | CCCTACCACTCGCCAATTCATCC |
| SD773_036340_qPCR R | GCAAGGGCGACAAATCCCG |
| SD375_h4_qpCR F | CCGCCGTGGTGGTGTCAAG |
| SD376_H4_qPCR R | GGCGTGTTCAGTGTAGGTGACG |
| SD537_insA_qPCR_New_F | GCCTCAGGTGTTCCGAAGCG |
| SD538_insA_qPCR_New_R | CCAGAGCCACTCCAGCCAGA |
| SD160_afu182_F_new | GCAGTCCCTTCCACCGAGC |
| SD161_afu182_R_new | GCGTTCCTACGAGGATTATCCTCC |
| SD539_srbA_qPCR_New_F | TCGAGGCGAGCTCTGGTGG |
| SD540_srbA_qPCR_New_R | GCTTCCGATGTTGGCGACGC |
| SD541_erg11a_qPCR_New_F | GCGGTGCTGACGGCAATCT |
| SD542_erg11a_qPCR_New_R | CGTTTCCCTGAACGCCCAGG |
| SD543_erg11b_qPCR_New_F | CGGCGTTCCAGGGACACAA |
| SD544_erg11b_qPCR_New_R | CGCAGCATCCCTTTTGCGGT |
| SD545_srbB_qPCR_New_F | ACGGATCTGTTTGCTCCGCC |
| SD546_srbB_qPCR_New_R | CAGGACTGGACGACCAACGA |
| SD547_atr_qPCR_New_F | GGC GGT ACT CTG TGT GAC TGC |
| SD548_atr_qPCR_New_R | GCGCGAGCATCACTGGGATTC |
| SD363_18s Forward | GGC CCT TAA ATA GCC CGG T |
| SD364_18s R | TGA GCC GAT AGT CCC CCT AA |
| SD365_18s probe | /56-FAM/AG CCA GCG G/ZEN/C CCG CAA ATG /3IABkFQ/ |

| | |
|---|---|
| SD843_afub_033480_qPCR_F | CATTCATCCTGGTCGGGTCT |
| SD490_afub_033480_qPCR_R | GCTCTCTTCGCTTCTTTCTTCTCGC |

Table S3. List of crRNAs used in this study

| Name | crRNA sequence |
|---|---|
| 60R_gRNA1 | AltR1/rGrU rCrArA rCrUrC rGrGrU rUrUrC rArGrA rArGrU rGrUrU rUrUrA rGrArG rCrUrA rUrGrC rU/AltR2/ |
| 60L_gRNA 2 | /AltR1/rArU rUrCrC rGrArG rUrGrA rArGrU rUrGrC rArCrG rGrUrU rUrUrA rGrArG rCrUrA rUrGrC rU/AltR2/ |
| 50L_gRNA3 | /AltR1/rCrC rUrUrU rCrCrA rGrGrC rGrUrA rArUrA rUrCrA rGrUrU rUrUrA rGrArG rCrUrA rUrGrC rU/AltR2/ |
| 50R_gRNA 4 | /AltR1/rUrC rUrGrC rUrCrA rArArC rCrGrG rGrCrA rArUrA rGrUrU rUrUrA rGrArG rCrUrA rUrGrC rU/AltR2/ |
| 182 L_gRNA5 | /AltR1/rCrU rUrCrG rUrCrC rArGrA rUrUrC rUrArC rArArA rGrUrU rUrUrA rGrArG rCrUrA rUrGrC rU/AltR2/ |
| 182 R_gRNA6 | /AltR1/rUrA rUrUrU rGrCrG rUrUrG rUrUrC rArUrG rArArG rGrUrU rUrUrA rGrArG rCrUrA rUrGrC rU/AltR2/ |
| gRNA 13 SD 1 | /AltR1/rGrG rUrCrG rCrCrU rCrArA rArCrA rArUrG rCrUrC rGrUrU rUrUrA rGrArG rCrUrA rUrGrC rU/AltR2/ |
| atf4_gRNA7 | /AltR1/rUrC rUrCrC rUrUrC rArUrA rArGrC rGrArC rCrArG rGrUrU rUrUrA rGrArG rCrUrA rUrGrC rU/AltR2/ |
| nidulans 182 F_gRNA8 | /AltR1/rArU rGrArU rUrGrG rUrUrC rUrArC rArGrA rCrUrU rGrUrU rUrUrA rGrArG rCrUrA rUrGrC rU/AltR2/ |
| nid 182 R_gRNA9 | /AltR1/rGrC rCrGrC rGrCrA rGrUrG rGrUrC rUrGrU rGrCrG rGrUrU rUrUrA rGrArG rCrUrA rUrGrC rU/AltR2/ |
